# Mapping the distribution of seagrass meadows from space with deep convolutional neural networks

**DOI:** 10.1101/2024.03.21.586047

**Authors:** Àlex Giménez-Romero, Dhafer Ferchichi, Pablo Moreno-Spiegelberg, Tomàs Sintes, Manuel A. Matías

## Abstract

Seagrass meadows play a vital role in supporting coastal communities by promoting biodiversity, mitigating coastal erosion and contributing to local economies. These ecosystems face significant threats, including habitat loss and degradation or climate change. United Nations has recognized the urgency of conserving marine ecosystems, highlighting the need for evidence-based conservation strategies and high-quality monitoring. However, traditional monitoring approaches are often time-consuming, labor-intensive, and costly, limiting their scalability and effectiveness. The growing availability of remote sensing data coupled to the rise of machine learning technologies offer an unprecedented opportunity to develop autonomous, efficient and scalable monitoring systems. Despite many efforts, the development of such systems for seagrass meadows remains a challenge, with recent attempts presenting several limitations such as limited satellite imagery, inadequate metrics for evaluating model performance or insufficient ground truth data, leading to simple proof of concepts rather than useful solutions. Here, we overcome these limitations by developing a comprehensive framework to map *Posidonia oceanica* meadows in the Mediterranean Sea using an extensive georeferenced habitat dataset and diverse satellite imagery for model training. We successfully evaluate the model generalization capability across different regions and provide the trained model for broader application in biodiversity monitoring and management.

---

Coastal ecosystems, encompassing seagrasses, mangroves, saltmarshes, and coral reefs, among others, provide invaluable services that contribute to support the livelihoods of coastal communities, impacting the well-being of the residents [1, 2]. Seagrass meadows, in particular, are crucial for enhancing coastal biodiversity by serving as essential habitats for a diverse range of marine species [3]. They provide vital food, shelter, and structural support, including nursery areas for commercially important species, thereby supporting both local economies and subsistence fisheries [4]. Moreover, seagrass ecosystems play a significant role in coastal erosion prevention. The dense canopies of seagrass attenuate currents and waves, facilitating particle sedimentation and mitigating sediment resuspension [5–8]. Additionally, the extensive underground network of rhizomes and roots stabilizes sediment, reducing erosion and decreasing water turbidity [9], strongly influencing coastal sedimentary dynamics [10, 11]. Specifically, the robust root systems of *Posidonia oceanica* beds act as natural barriers, protecting coastlines from the destructive force of strong waves and maintaining shoreline stability [12–14]. In addition, seagrasses keep overlying waters oxygenated and with low concentrations of nutrients and CO_2_ [15]. Seagrasses are among the planet’s most effective natural ecosystems for sequestering (capturing and storing) carbon, performing a rate that is 35 times faster than tropical rainforests, while their sediments never become saturated [16].

However, if these habitats are degraded, they could leak stored carbon into the atmosphere and further accelerate global warming [17, 18]. In fact, despite the considerable uncertainty surrounding global seagrass extent values, it is estimated that about one third of seagrass global extent has been lost since World War II [17]. Seagrass declines are primarily attributed to eutrophication, water quality degradation, habitat destruction, and climate change, particularly global warming [19]. Furthermore, the sensitivity of seagrasses to future ocean temperatures under different emission scenarios poses significant concerns. Models project a decline in the global suitable habitat for these ecosystems throughout the current century, both latitudinally and across water depth, with a notable compression of suitable habitat toward the lower distribution limit imposed by light availability [20].

In this context, the United Nations (UN) recognized the severity of global biodiversity loss and degradation of ecosystems and stressed the negative impact that this situation has on food security, nutrition, access to water, health of the rural poor and people worldwide. Accordingly, the UN declared the period 2021-2030 as the “Decade of Ocean Science for Sustainable Development” and the “Decade of Ecosystem Restoration” [21, 22], underlining the urgency and importance of safeguarding marine ecosystems, including *Posidonia oceanica* meadows. Achieving these targets, particularly concerning the preservation and restoration of coastal ecosystems, requires a rigorous, evidence-based approach to conservation practice and policy. This entails conducting thorough analyses of high-quality monitoring data to inform decision-making and validate intervention strategies.

The comprehensive mapping of several marine habitats, such as coral reefs, kelp forests, deep-sea vent communities, and seagrass beds, has been successfully achieved through the use of sidescan sonar systems [23–26]. This methodology has provided valuable insights into the structure and distribution of these ecosystems and helped to design informed conservation strategies and management practices. However, the cost and time-intensive nature of these methods present challenges in deploying continuous monitoring systems for marine environments. As a result, practical monitoring of biodiversity often occurs infrequently rather than in real-time, preventing a constant spatiotemporal evaluation of the status of these ecosystems.

A recently emerging possibility is to combine remote sensing technologies with available georeferenced habitat data to develop correlative or mechanistic models, which are then capable to monitor biodiversity at finer temporal scales. Among various methodologies, Machine Learning (ML) models trained with multi-spectral satellite imagery data appear to be the most promising [27–30]. In particular, many efforts to determine the spatial distribution of *Posidonia oceanica* from airborne imagery using ML have been recently made [31–48]. However, despite the seminal insights of many of these works, they present numerous limitations that hinder the delivery of functional models suitable for a real-case deployment, serving merely as potent proof of concept for the methodologies studied. For instance, many of these studies rely on inadequate metrics for evaluating model performance in image segmentation problems, such as accuracy, leading to an overestimation of the model’s performance and neglecting more suitable and demanding metrics like Intersection over Union. Additionally, ground truth data predominantly rely on photo-interpretation, often with a limited number of validation points obtained from field data, undermining the models’ robustness. Several studies employ only a single or few satellite images, limiting the models’ generalizability, while there’s a prevalence of simplistic ML methods like Supported Vector Machines and Random Forests for image segmentation tasks, despite the suitability of Deep Convolutional Neural Networks (CNN) for such tasks being well-established [49, 50]. Thus, while these studies lay the groundwork for innovative methodologies, further research is imperative to develop robust and scalable models capable of meeting the demands of real-world applications in biodiversity monitoring and management.

To reach this goal, three key considerations need to be addressed. Firstly, an extensive georeferenced habitat dataset must be employed, acquired through a meticulous and consistent methodology. This dataset should cover a broad geographical area and encompass various spatial scales to ensure the representation of diverse ecological conditions. Secondly, deep learning models, preferably based on convolutional neural networks (CNNs), should be trained using a diverse set of satellite images. These images should incorporate variations in acquisition dates, geographic locations, and the positions of satellites relative to Earth and the sun. This approach enables learning under real-world conditions and enhances the robustness of the models. Lastly, the generalization capability of the models must be evaluated by testing their predictive performance across regions that are geographically distinct from the training dataset. These regions should be characterized by different environmental conditions, ensuring the reliability and applicability of the models in varied real-world scenarios.

Here, we present a comprehensive framework that addresses these considerations, providing a robust and reliable model for the classification of *Posidonia oceanica* meadows and related habitats in the Mediterranean Sea using satellite imagery. We demonstrate the model’s generalization capability and robustness by training only with data from a particular region of our extensive dataset and evaluating its performance on the other regions. We show that our model is capable of providing reliable estimates for the distribution of the considered habitats and accurate measures for their extension area. In addition, we measure the model’s loss of accuracy in its estimates for new regions, providing a lower bound for the model performance. This is a crucial step to advance in the development of a reliable map of the distribution of *Posidonia oceanica* meadows in the Mediterranean Sea. Finally, we train the model with all available data so that the resulting model can be used to classify Posidonia meadows in other regions of the Mediterranean Sea.

## Results

### A deep learning framework for automated marine ecosystem labelling

We developed a deep learning framework based on convolutional neural networks to accurately classify benthic habitats in the Mediterranean Sea using satellite imagery (Fig. 1). We used a comprehensive and extensive habitat dataset of the Balearic sea, comprising a 20 year effort of data acquisition based on side-scan sonar supported by photo-interpretation of high-resolution airborne imagery and in-situ observations (Fig. 1a). The dataset covers about 2,500 km^2^ of the coastal habitats of the Balearic Islands at high spatial resolution and contains 28 different classes, including the ecologically significant species *Posidonia oceanica*, which were aggregated in 4 major ecological groups: Posidonia oceanica, Other green plants, Rocks & brown algae and Sandy bottoms (Fig. 1a,c, Methods & Supplementary Information). This dataset was combined with satellite imagery of the coastal areas of the Balearic Islands acquired from PlanetScope [51], covering around 1200 km^2^ with different dates and satellite positions (Fig. 1b,c, Methods & Supplementary Information).

**Figure 1:**
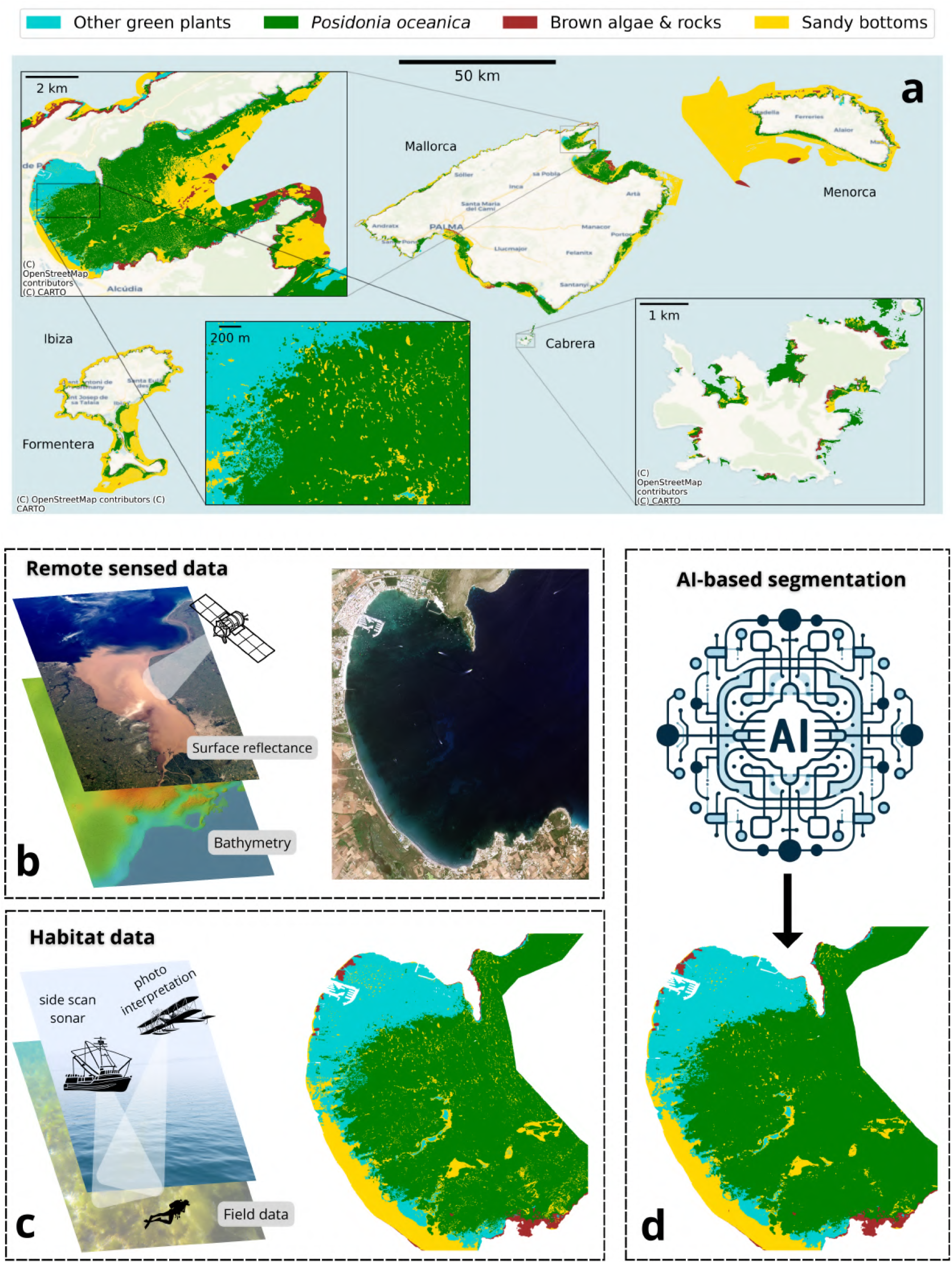
(a) Spatial distribution of the 4 main ecological benthic habitats in the Balearic Sea, present in the whole Mediterranean. The dataset provides detailed information at multiple spatial scales up to 3 m resolution. (b-d) Scheme of the pipeline to train CAMELE model. (b) Satellite-based surface reflectance data is merged with bathymetry estimates to produce the inputs (features) of the model. Habitat data obtained with side-scan sonar, photo-interpretation and field observations were used as ground truth data (labels). (d) These features and labels are used to train deep convolutional neural networks to perform image segmentation.

We trained 40 different deep learning models using 4 different state-of-the-art architectures and 10 different backbones for each architecture (Fig. 1d, Methods & Supplementary Information). Furthermore, we implemented a consensus prediction approach to enhance the robustness and reliability of model predictions, which involves aggregating the results from multiple deep learning models to mitigate potential biases introduced by individual models (Methods & Supplementary Information). To evaluate the models, we opted for training only with data from one island (Mallorca), performing a posterior systematic study of its performance on the other islands (Menorca, Ibiza, Formentera, and Cabrera). Thus, the train-test split was roughly 50%-50% rather than the traditional 80%-20% split, with the test set representing diverse environmental conditions and benthic habitats formed by slightly different species than the training set (Methods & Supplementary Information). This approach was chosen to simulate real-world scenarios, in which one cannot control for specific environmental conditions, constrained dates for image acquisition, the position of the satellite with respect to the sun and the earth, or even find new species not contained in the original training dataset. We thereafter refer to our test set as “out-of-sample” test set and to the model as “Half model” to emphasize this idea.

We performed and extensive evaluation of the models’ performance in the training and out-of-sample test datasets, using a variety of metrics such as Intersection over Union (IoU), Precision, Recall, F1-score, Kappa and Accuracy (Methods). Our results show that the best performing framework was to use the 10 models defined by the Linknet architecture together with the consensus prediction approach (Methods and Supplementary Information), which hereafter we refer to as CAMELE (Consensus for Automated Marine Ecosystem Labelling and Evaluation).

### A reliable AI-based solution for marine ecosystem monitoring

CAMELE’s performance in both training and out-of-sample test datasets was highly notable, with a mean IoU score of 88.22% and a mean F1-score of 93.13% in the training dataset, compared with a mean IoU score of 61.97% and a mean F1-score of 72.77% in the out-of-sample test dataset (Fig. 2a, and Supplementary Tables 5 and 6). We note that in image segmentation tasks an IoU score greater than 50% is considered an acceptable prediction [52] (yellow stars in Fig. 2a). Furthermore, the model outperforms by a large margin the naive baseline of predicting only the majority class (yellow diamonds in Fig. 2a). We observe an overlap between the distribution of the performance metrics in the training and out-of-sample test datasets, showing that model performance is consistent in both sets (Fig. 2a). The model was able to segment some images in the out-of-sample test dataset with notable performance (e.g. 15% of the images with an IoU score higher than 80%, and 20% of the images with an IoU score higher than 70%), while only 10% of the images had an IoU score lower than 50% (Fig. 2a and Supplementary Table 8). This demonstrates that the model is able to generalize to some extent to new regions, with different environmental conditions and the presence of some different benthic habitats.

**Figure 2:**
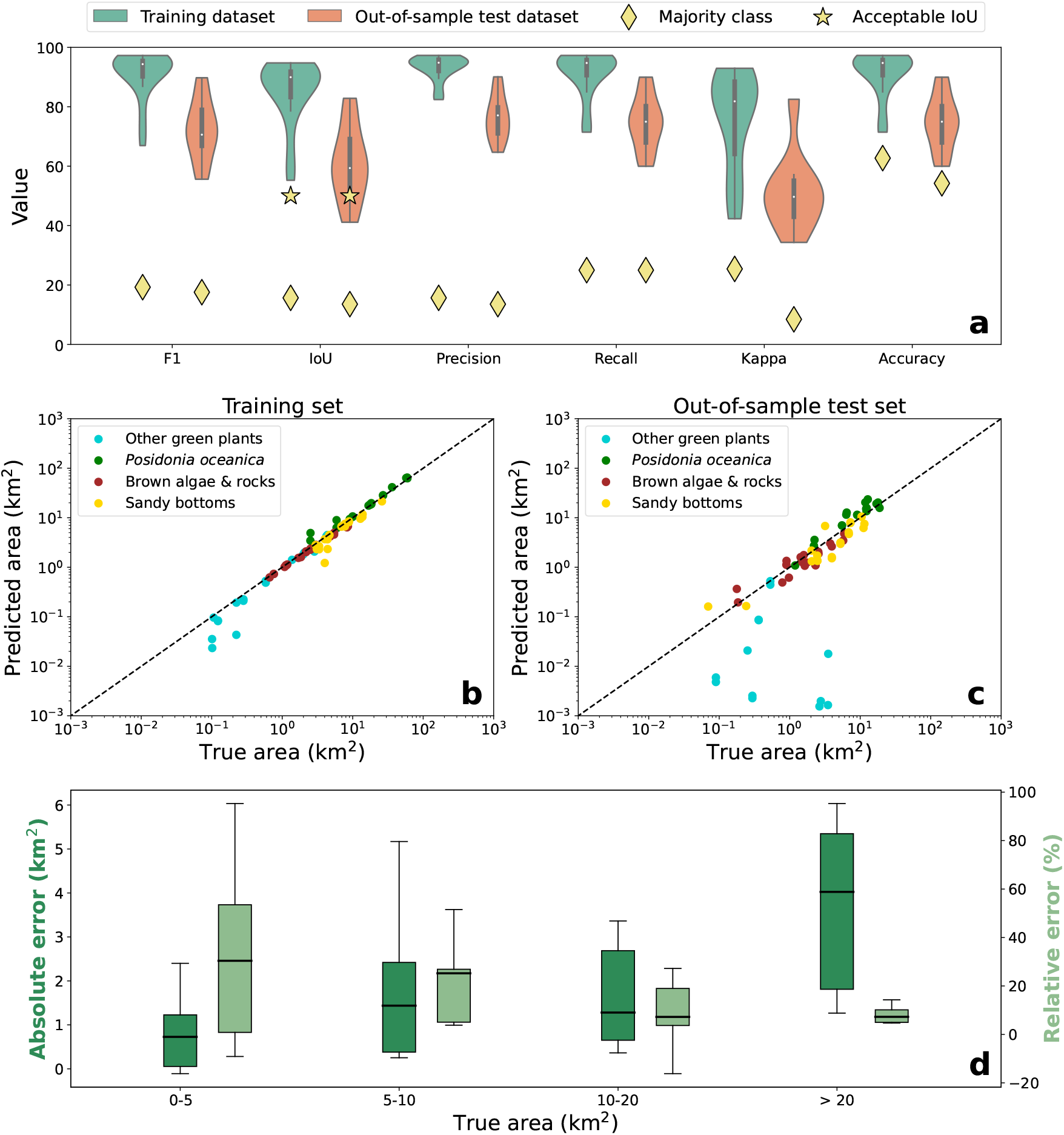
Model performance in train and out-of-sample test datasets. (a) Violin plots for F1-score, IoU, Precision, Recall, Kappa and Accuracy in both training and out-of-sample test datasets. (b-c) True vs predicted area for each habitat class in the training (b) and out-of-sample test (c) datasets. The diagonal dashed line indicate perfect prediction. (d) Box plot for the absolute and relative errors commited in the prediction of *Posidonia oceanica* area as function of the true area. Relative errors significantly decrease with the extent of the area to be predicted, linked to the wider spatial context available.

Despite the decrease in performance in the out-of-sample test dataset can be attributed to the different environmental conditions, including the presence of some different benthic habitats, we observed that a significant part of the pixels categorized by the ground truth data as Other green plants, Brown algae & rocks or Sandy bottoms were being classified by the model as Posidonia oceanica, substantially affecting the overall performance (Supplemantary Information, Supplementary Fig. 3). Surprisingly, we found that the distribution of response values for the Other green plants class in the test dataset is much more similar to the distribution of response values of Posidonia oceanica class in the train set than to its own class (Supplementary Information, Supplementary Fig. 4). Then it is not surprising that the model classifies all those samples as *Posidonia oceanica*. In contrast, the model achieved notable performance in segmenting the Posidonia oceanica class, with a mean IoU of 77.30% compared with the mean IoU of 91.97% achieved in the training dataset (Supplementary Table 7). At any rate, the model still achieved an overall notable performance in the out-of-sample test dataset, demonstrating the generalization capability and robustness of CAMELE and highlighting its potential for real-world applications in biodiversity monitoring and management.

CAMELE’s performance was further evaluated by comparing the true and predicted area for each habitat class in each image of the training and out-of-sample test datasets. The model achieved a notable performance in predicting the area of the different habitat classes except for the Other green plants class, as expected from the previous analysis (Fig. 2b,c). Specifically, the median absolute errors committed in the prediction of the area of the different habitat classes were 0.98 km^2^, 0.06 km^2^, 0.15 km^2^ and 0.70 km^2^ in the training dataset, compared with 2.24 km^2^, 0.26 km^2^, 0.47 km^2^ and 1.12 km^2^ in the out-of-sample test dataset, for the Posidonia oceanica, Other green plants, Rocks & brown algae and Sandy bottoms classes, respectively. However, the relative errors were 5.61%, 11.77%, 6.77% and 14.34% in the training dataset, compared with 24.92%, 99.20%, 28.16% and 35.05% in the out-of-sample test dataset. Of course, the high relative errors for the Other green plants class in the out-of-sample test dataset are nonsensical, as the true area of this class is small, leading to a high relative error. Thus, we observe that the absolute errors doubled in the out-of-sample test dataset, while the relative errors increased by a factor of 5. At any rate, the models’ performance in predicting the area of the Posidonia oceanica class was particularly notable, with relative errors significantly decreasing with the extent of the area to be predicted, linked to the wider spatial context available (Fig. 2d). For instance, the median relative error for true extent areas between 1 and 5 km^2^ was 30% (86% for 95% confidence interval) compared with 7% (86% for 95% confidence interval) for areas larger than 20 km^2^. This finding underscores the importance of considering the spatial context when predicting the area of benthic habitats, highlighting the potential of CAMELE to provide reliable estimates for the distribution and extension of the considered habitats in the Mediterranean Sea.

To further illustrate the model’s performance, we present an example of model predictions for a satellite image in the training and out-of-sample test sets (Fig. 3). The model accurately classified the different benthic habitats in both cases with 92.78% and 82.54% IoU scores, respectively, providing reliable estimates for the distribution and extension of the considered habitats. We note that although there is a 10 point difference in the IoU score between each prediction, this is almost unobservable in the visual inspection of the predictions, highlighting the extreme sensitivity of this metric to small differences.

**Figure 3:**
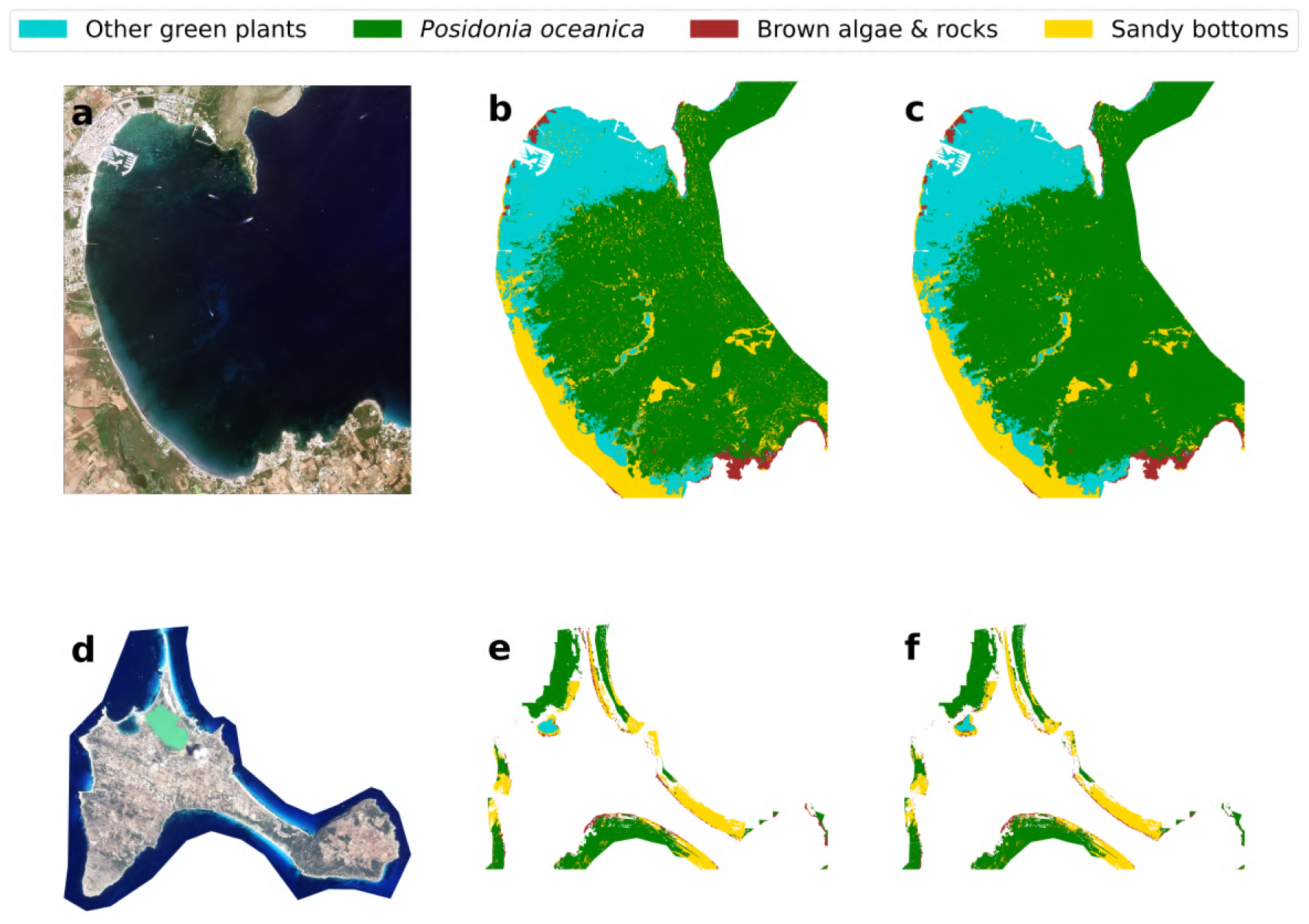
Example of model predictions for a satellite image in the training and out-of-sample test set. (a) Satellite image from Pollença bay in the island of Mallorca, a part of the training set. Image © 2022 Planet Labs PBC (b) Ground truth data for the benthic habitats in Pollença bay. (c) Habitat classification from CAMELE model in Pollença bay (92.78% IoU). (d) Satellite image from Formentera island, part of the out-of-sample test set. Image © 2022 Planet Labs PBC (e) Ground truth data for the benthic habitats in Formentera. (f) Habitat classification from CAMELE model in Formentera (82.54% IoU).

### Towards a comprehensive model for the Mediterranean Sea

Finally, we trained CAMELE with all available data (using 13144 patches for the actual training and 3286 for the validation set) and provide the scientific community with our final trained models, which are freely accessible at [53]. We thereafter refer to this model as the “final” model. We evaluated the performance of the final model in the complete dataset and, for comparison, also in the previous training and out-of-sample test datasets, achieving a median IoU score of 95.22%, 94.73% and 96.22%, respectively (Table 1). Notably, the model’s segmented all the images in the complete dataset with an IoU score higher than 90%, with a mean, median and maximum IoU score of 94.64% 95.22% and 98.5%, respectively (Supplementary Table 10). Finally, we assessed the model robustness by predicting on a new set of images from the Balearic Islands in 2023, not used in the training or out-of-sample test datasets. The model achieved a remarkable mean IoU score of 80% in this new dataset (Supplementary Table 11).

**Table 1:** Performance (IoU score) of the final model in the complete dataset and comparison with the previous model performance and previous data splits conforming the training and out-of-sample (OOS) test datasets.

|  | Train (Half Model) | OOS test (Half Model) | Complete dataset |
| --- | --- | --- | --- |
| **Half model** | 88.22 | 61.97 | 79.21 |
| **Final model** | 94.73 | 96.22 | 95.22 |

In Fig. 4, we present some examples of model predictions in different regions of the Balearic Islands, showing the ground-truth data for the benthic habitats, the habitat classification from CAMELE model trained only with half of the data (from Mallorca island), and the habitat classification from CAMELE model trained with all available data. We observe that the main differences between the predictions of the two models occur in the areas with more complex habitat distribution. In these areas, the model trained with all available data is able to capture the complexity of the habitat distribution more accurately, providing a more detailed and reliable classification of the benthic habitats. In any case, the model trained only with data from Mallorca still provides a notable performance in segmenting the *Posidonia oceanica* meadows from the other islands, which in the end is the most important habitat to be monitored. A web-based application for interactively visualizing all model’s predictions, together with the ground truth data, is available at [54].

**Figure 4:**
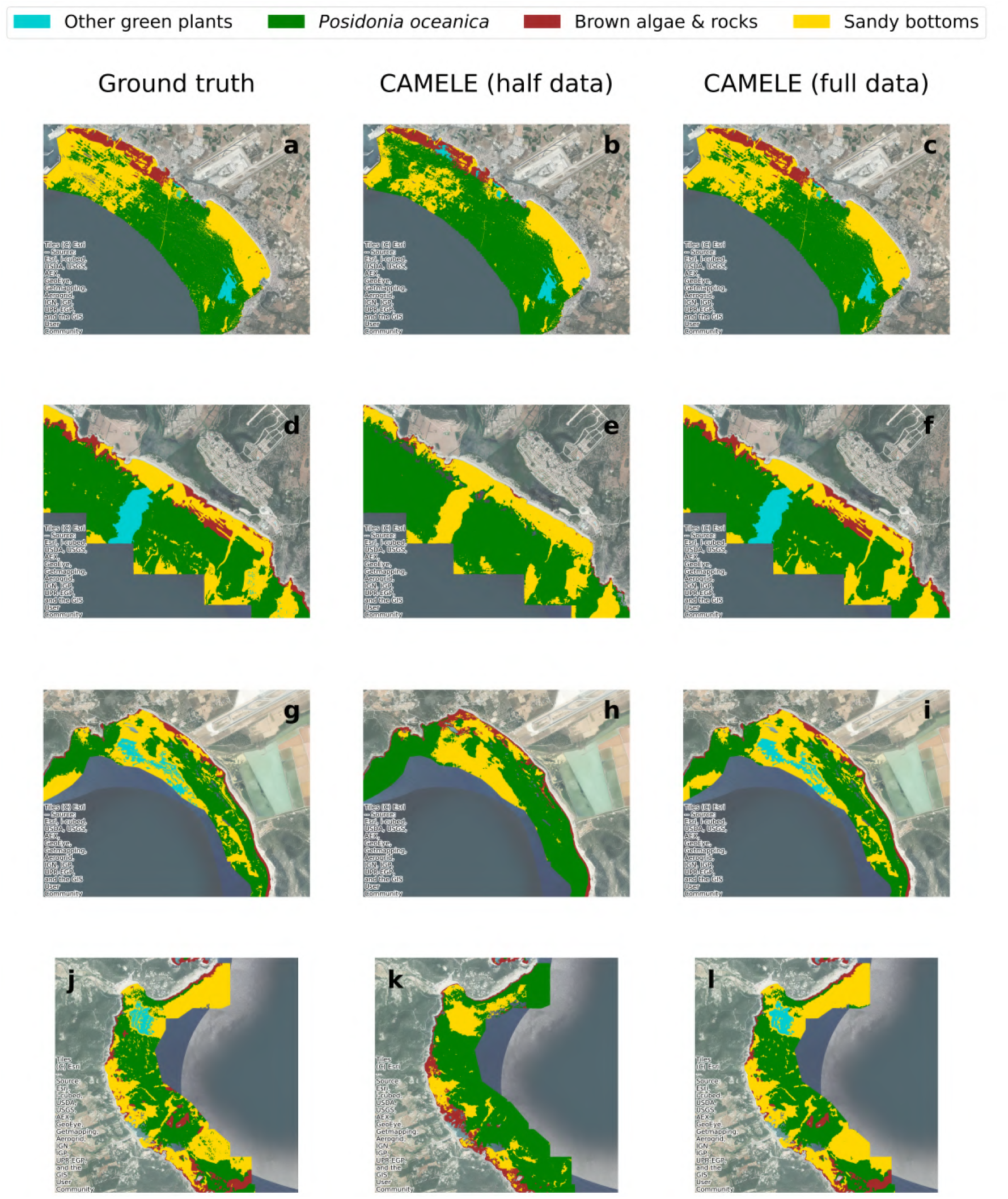
Example of model predictions for a satellite image in the complete dataset. The first column shows the ground truth data for the benthic habitats, the second column shows the habitat classification from CAMELE model trained only with half of the data (from Mallorca island), and the third column shows the habitat classification from CAMELE model trained with all available data. (a-c) Palma bay, Mallorca. (d-f) Son Bou beach, south east of Menorca. (g-i) Es Còdols, south of Ibiza. (j-l) Cala San Vicente, east of Ibiza.

## Discussion

The present study represents a significant advancement in the field of marine habitat monitoring, leveraging the synergistic combination of remote sensing data and machine learning algorithms to address critical challenges in biodiversity conservation. Unlike previous studies, we have developed a comprehensive framework for classifying *Posidonia oceanica* meadows from multi-spectral satellite imagery based on deep convolutional neural networks, which in the last years have been shown to be highly effective in image segmentation tasks. This approach takes advantage of both the rich spectral and spatial information provided by satellite imagery, and the capacity of deep learning models to learn complex patterns and relationships in the data. In addition, our analysis do not rely on a single or few satellite images, but rather on a comprehensive dataset of satellite images, covering a wide geographical area and various spatial scales, to ensure representation of diverse ecological conditions. This approach is crucial for training models under real-world conditions and enhancing their robustness and generalization capability. Indeed, here we make a substantial effort to evaluate the model’s generalization capability by testing its predictive performance across regions geographically distinct from the training dataset, characterized by different environmental conditions, to ensure its reliability and applicability in varied scenarios. Of course, this would not have been possible without the extensive and detailed georeferenced habitat dataset used in this study.

One of the key findings of our study is the remarkable generalization ability of our deep learning model across different regions of the Balearic Islands. Despite being trained exclusively on data from a specific area, the model demonstrated the capacity to provide reliable estimates of *Posidonia oceanica* distribution in other islands, where environmental conditions and benthic habitats vary. This highlights the robustness of our approach and its potential for broader applicability in marine habitat monitoring. The ability of the model to generalize across regions is crucial for its practical utility in conservation efforts beyond the training domain. By accurately classifying marine habitats in unseen environments, our model showcases its capacity for real-world application, particularly in scenarios where comprehensive training data may be limited. Furthermore, the provided metrics, including median absolute and relative errors in *Posidonia oceanica* area prediction, offer valuable insights into the model’s performance and potential limitations. By quantifying prediction errors and their relationship to true area estimates, we establish a foundational understanding of the model’s accuracy and reliability, enabling more informed decision-making in habitat monitoring and conservation efforts.

The combination of remote sensing and Machine Learning holds promise for revolutionizing biodiversity monitoring, offering unprecedented opportunities for conservationists, policymakers, and researchers to make informed decisions and address pressing environmental challenges. With its ability to provide near-real-time data on ecosystem dynamics, our approach offers a cost-efficient and scalable solution for continuously assessing biodiversity distribution in the Mediterranean sea, allowing to identify habitat degradation and monitor ecosystem resilience in the face of environmental change. This is particularly crucial in the context of the ongoing climate crisis, where the ability to rapidly detect and respond to habitat loss and degradation is essential for preserving marine biodiversity and ecosystem services. Thus, our model represents a powerful tool in the conservation toolbox, enabling timely interventions supporting the sustainable management of coastal ecosystems and the preservation of biodiversity for future generations.

Despite the notable advancements and potential of our study, several challenges and limitations remain to be addressed. Firstly, the reliance on satellite imagery for habitat classification presents inherent limitations, including cloud cover, atmospheric interference, and limited spatial and spectral resolution, which may impact the accuracy and detail of habitat classification, especially in complex coastal environments. Additionally, the availability and quality of training data pose challenges, as incomplete or biased datasets can impact model performance and generalization capabilities. Moreover, our analysis focused on 4 major aggregated ecological groups, including *Posidonia oceanica* habitats, neglecting other important benthic species and ecosystems that contribute to overall marine biodiversity. Furthermore, while our models exhibit robustness in cross-regional generalization within the Balearic Islands, their applicability to other geographical regions with distinct environmental conditions remains untested. Thus, it may exhibit limitations in delineating finer-scale habitat features in regions with distinct ecological characteristics. Addressing these limitations through continued data collection, model refinement, and validation efforts will be crucial for advancing the reliability and applicability of remote sensing-based approaches in marine habitat monitoring and conservation.

Looking ahead, several avenues for future research and practical applications emerge from our study. Firstly, continued efforts to expand and refine training datasets, incorporating data from diverse geographic regions and ecosystem types, will further enhance the accuracy and generalization capabilities of machine learning models for marine habitat monitoring. Additionally, ongoing advancements in remote sensing technology, including the development of higher spatial and spectral resolution sensors, hold promise for improving habitat classification accuracy and detail. Integration with emerging techniques such as drone-based imaging and LiDAR further expands the scope and resolution of habitat monitoring efforts, enabling finer-scale analysis and management. Moreover, the development of user-friendly tools and platforms for data visualization and decision support facilitates the translation of scientific findings into actionable conservation strategies. By empowering stakeholders with accessible and interpretable information, we can foster greater engagement and collaboration in marine conservation initiatives, ultimately contributing to the sustainable management of coastal ecosystems and the preservation of biodiversity.

In conclusion, our study represents a significant step forward in the field of marine habitat monitoring, showcasing the transformative potential of remote sensing and machine learning technologies. By advancing our understanding of ecosystem dynamics and supporting evidence-based conservation practices, our work contributes to the broader mission of safeguarding marine biodiversity for future generations.

## Methods

### Satellite data

Satellite imagery was obtained from Planet under the Education and Research Program, which provides limited, non-commercial access to PlanetScope and RapidEye imagery [51]. In particular, we adcquired PlanetScope images obtained through the Super Dove (PSB.SD) instrument, which consists of Coastal Blue, Blue, Green I, Green, Yellow, Red, Red Edge, NIR and operates from 2020. Surface Reflectance (SR) products were selected to ensure consistency across localized atmospheric conditions, minimizing uncertainty in spectral response across time and location. SR is derived from the standard Analytic Product (Radiance) and is processed to top-of-atmosphere reflectance and then atmospherically corrected to bottom-of-atmosphere reflectance.

We obtained a total of 60 satellite images along the coast of the Balearic Islands, covering up to 1200 km^2^ surface area for the years 2020 to 2023 (see supplementary information, Supplementary Fig. 1). The images were acquired for days with clear sky conditions comprised between June and September, as these are the months in which the biomass of seagrass and algae is more abundant. No other filters were applied to the images, as we aimed to train the model in real-case scenarios, in which one can not control for specific environmental conditions, constrained dates for image acquisition, the position of the satellite with respect the sun and the earth, etc. Thus we obtained images with different conditions (see Supplementary Information, Supplementary Table 1).

### Habitat data

We used georeferenced habitat data from the government of the Balearic Islands, corresponding to the outcome of different European and national projects comprising a *∼* 20 year effort of data acquisition covering a total of 2,500 km^2^ [55, 56]. The first project started around the year 2000 and there has been a major recen update around 2018. The data consist of 28 different habitat classes following the nomenclature and coding of the Standard List of Marine Habitats of Spain (LPHME), obtained from side scan sonar, photo-interpretation of airborne imagery and in-situ observations. We aggregated the different habitat classes in 4 major ecological groups, present in the whole Mediterranean sea. The aggregation was based on feature similarity and ecological function, resulting in the following classes: Posidonia oceanica, green algae, brown algae & rocks and sandy bottoms (see Supplementary Information, Supplementary Table 2).

Fig. 1a shows the spatial distribution of the considered ecological groups in the Balearic Sea. This comprehensive dataset provides detailed information at multiple spatial scales, offering a valuable insight into the intricate spatial patterns of the different habitat classes across the region. The detailed information captured by the high-resolution data contributes significantly to the reliability of our model and the precision of our predictions, particularly vital in the context of habitat conservation for *Posidonia oceanica* meadows.

### Bathymetry data

Bathymetry data was obtained from the European Marine Observation and Data Network (EMODNET) [57]. EMODnet-Bathymetry provides a service for viewing and downloading a harmonised Digital Terrain Model (DTM) for the European sea regions. The data consist of GeoTIFF layers with *∼* 100 m pixel resolution of mean depth values.

### Dataset creation

Satellite images were processed together with habitat and bathymetry data to construct the final training and testing datasets. The NIR band, which is strongly attenuated by water, was used to mask out pixels corresponding to land using a simple clustering algorithm (i.e. K-means) and subsequently substituted for bathymetry data. The resulting processed satellite images comprise the primary source for the input data of our model. The ground truth (or label) dataset consist of raster files analogous to the satellite image with single-band values representing the benthic class corresponding to each pixel. To construct it, all pixels of each processed satellite image were associated to a given benthic class using the aggregated habitat data. Pixels for which a class was not available were masked out in both the processed satellite image and label data. Similarly, pixels already masked in the satellite image were masked in the label image. Finally, patches measuring 256 x 256 pixels (*∼* 750 *m* ***×*** 750 *m*) were extracted from each satellite and label image for the years 2020 to 2022 (keeping the year 2023 as a final test), resulting in a final dataset comprising up to 16,430 patches.

Despite train-test data split usually follows a 80%-20% ratio, training was conducted exclusively with data from Mallorca (8488 patches, of which 1698 were allocated for validation) and tested with the remaining data from Menorca, Ibiza, Formentera, and Cabrera islands (7942 patches), following a roughly 50%-50% train-test split. Furthermore, our test set represented diverse environmental conditions and even benthic habitats formed by slightly different species than the training set, thereby simulating real-world scenarios that the model may encounter in operational settings. By testing the model on data from regions beyond its training domain, we aimed to assess its generalization ability and robustness. Specifically, we sought to determine whether the model could accurately classify marine habitats and benthic features in unseen environments, probably slightly different from the ones at training, thereby demonstrating its capacity for real-world application. We thereafter refer to our test set as “out-of-sample” test set, to emphasize this idea. All images from 2023 were kept for a final test to analyze model robustness once it is trained with data from all regions in the years 2020 to 2022.

### Deep learning models

We trained different state-of-the-art deep learning models for semantic image segmentation, such as UNET [58], Linknet [59], FPN [60], and PSPNet [61]. Each architecture is formed by repeating convolutional blocks, which are usually referred to as the “backbone” of the model. We tested 10 different backbone models for each architecture, leading to the training and evaluation of 40 deep learning models, which represents and unprecedented effort in the field. We used the segmentation-models Python library [62] to define and train all models.

### Model training

Before training, the input data was standardized using the mean and variance of the training data,

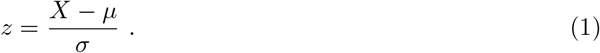

This standard scaling procedure ensures that all input features have a consistent scale, preventing the dominance of certain features during the training process. It is crucial to note that the same standard scaling procedure must be applied during predictions, using the mean and standard deviation computed from the training set. This consistency ensures that the model interprets new data in a comparable manner to the training data.

Additionally, for the categorical nature of the output, labels were one-hot encoded. This encoding converts categorical labels into binary vectors, where each class is represented by a unique binary value, facilitating the model’s interpretation of the multi-class classification task.

In terms of loss function selection, we opted for dice loss due to its effectiveness in handling imbalanced datasets, a common characteristic in tasks involving semantic segmentation [63]. Dice loss, also known as the Sørensen–Dice coefficient [64, 65], measures the similarity between predicted and ground truth data by computing the intersection over the union of the two. To further account for class imbalance, we applied loss weights inversely proportional to the proportion of examples of each class. Learning rate was set to 0.001 to ensure an smoother training process.

The models were trained in a computing cluster, using 10 cores and a maximum of 400 GB of RAM for each model. The training process was performed for 1000 epochs with a batch size of 32. The total training time was approximately 1 month for the 40 initial models using the data from the island of Mallorca and about 3 months for the 10 final models using all available data.

### Performance metrics

The performance of our trained models was primarily evaluated by the Intersection over Union (IoU) score (Eq. (7)), which evaluates the spatial overlap between the predicted image and the ground truth, as this is a suitable metric for image segmentation problems [63]. Additionally, we considered other metrics such as accuracy (Eq. (2)), precision (Eq. (3)), recall (Eq. (4)), F-1 score (Eq. (5)) and Cohen’s Kappa (Eq. (6)) to perform a comprehensive evaluation of the model. Accuracy gauges the overall correctness of predictions, precision measures the accuracy of positive predictions, recall assesses the model’s ability to capture all positive instances, and the F-1 score provides a balanced assessment of precision and recall. For a binary classification problem, the metrics can be define from the confusion matrix, where TP, TN, FP and FN represent true positives, true negatives, false positives and false negatives, respectively, and *N* is the total number of pixels in the image.

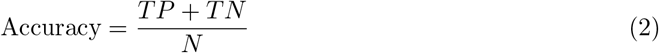

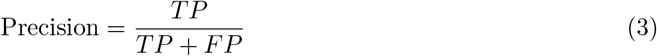

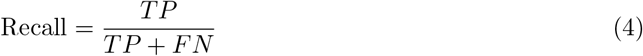

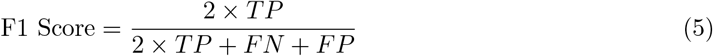

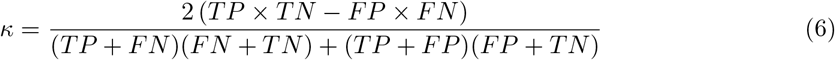

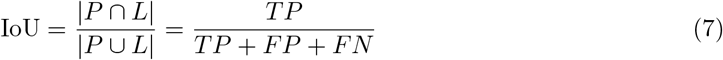

These metrics collectively offer a holistic understanding of the model’s effectiveness in classifying benthic habitats. See Supplemantary information for further details.

### Consensus prediction

We implemented a consensus prediction approach to enhance the robustness and reliability of model predictions. The consensus prediction involves aggregating the results from multiple deep learning models, each trained with different architectures and backbones. By combining predictions from diverse models, we aim to mitigate potential biases introduced by individual models and enhance the overall accuracy and generalization capabilities.

For each input patch, predictions from all trained models were collected, and a voting mechanism was employed to determine the final consensus prediction. Specifically, the class label with the highest frequency across all model predictions was assigned to the each pixel. This ensemble-based strategy leverages the diversity of information captured by different models, leading to a more robust and reliable classification outcome.

### Model selection

To filter among the 4 different architectures, we evaluated the models’ performance in the training dataset. Despite all models achieved high IoU scores (*>* 0.8), Linknet and UNET architectures were the best performing models with a mean IoU of 90.98% and 90.90%, respectively (Supplementary Table 3). Because Linknet architecture has less trainable parameters than UNET, being more efficient in terms of computational resources and less prone to overfitting, we selected it as the final architecture to build CAMELE, finally consisting of 10 different models (Supplementary Information).

We then evaluated the performance of each of the 10 models and the consensus prediction approach in the training and out-of-sample test datasets. All models achieved high performance, with a median IoU of 88.12% and F1-score of 93.16% in the training dataset (Supplementary Table 5), while the models’ performance in the out-of-sample test dataset was significantly reduced, with a median IoU of 60.73% and F1-score of 71.87% (Supplementary Table 6). When analyzing the performance of the models individually, we observed the emergence of “specialists”, which performed significantly better than the rest of the models in segmenting specific classes. This finding underscores the importance of the consensus prediction approach, which leverages the diversity of information captured by different models. The consensus prediction approach significantly improved the models’ performance in the out-of-sample test dataset, with an IoU score of 61.97% and a F1-score of 72.77% (Supplementary Table 6), highlighting the effectiveness of the ensemble-based strategy in mitigating potential biases introduced by individual models and enhancing the overall accuracy and generalization capabilities of CAMELE. Thus, the consensus prediction approach was selected as the final model for CAMELE.

## CRediT authorship contribution statement

AGR, TS and MAM conceptualised the project; AGR, DF and PMS conducted investigations; AGR and DF processed and curated the data; AGR and DF developed the methodology and trained the models; AGR developed the web-based application; AGR wrote the original draft; AGR, DF, PMS, TS and MAM reviewed and edited the manuscript; TS and MAM supervised the project; TS and MAM acquired funding.

## Acknowledgements

This work was supported through grants TED2021-131836B-I00 (SEDIMENT) funded by Spanish Ministry of Science and Innovation MICIU/AEI/10.13039/501100011033 and by the European Union NextGenerationEU/PRTR Program; PID2021-123723OB-C22 (CYCLE) funded by MICIU/AEI/10.13039/501100011033 and by “ERDF A way of making Europe”; CEX2021-001164-M (María de Maeztu Program for Units of Excellence in R&D) funded by MICIU/AEI/10.13039/501100011033. We acknowledge Nuria Marbà for her valuable initial comments and suggestions. We acknowledge Marcial Bardolet and Jorge Moreno, from the Species Protection Service of the Government of the Balearic Islands, to provide the habitat data used in this study. We express our heartfelt gratitude to José Luis Cantero Rada for being a profound source of inspiration.

## Data and code availability

The final full-trained deep learning models employed to build CAMELE are available at [53]. Because satellite imagery was obtained from Planet under the Education and Research Program, which provides limited, non-commercial access to PlanetScope and RapidEye imagery, we are unable to share the raw data. However, we provide two processed images together with the ground truth data and a code example to perform predictions with the final trained models [53]. The web-based application for interactively visualizing all model’s predictions, together with the ground truth data, is available at [54].

