## Supplementary Information for "Mapping the distribution of seagrass meadows from space with deep convolutional neural networks"

#### Contents

**Satellite imagery and ground truth data** 2

**Dataset creation** 5

**Deep learning models** 5

**Performance Metrics** 7

**Spectral reflectance analysis** 8

**Architecture selection** 9

**Out-of-sample predicting power and robustness** 10

**Effect of depth on model performance** 16

**CAMELE trained with all available data** 17

### Satellite imagery and ground truth data

#### Study region covering

We obtained a total of 60 Planetscope satellite images covering the coast of the Balearic Islands for the years 2020 to 2023. From those, the 20 images covering the island of Mallorca for years 2020-2022 were used to conform the training dataset, while the images from the other islands for the years 2020-2022 were included in the out-of-sample test set (Fig. 1). The resting images from the whole region in the year 2023 were lft as final test. The metadata of the satellite images used in the study is shown in Table 1.

Here we must emphasize that we refer to this test set as “out-of-sample” because the images conforming the set are relatively distant to the ones in the training set. Furthermore, the sub-classes conforming the 4 major ecological classes are not exactly the same in each island (see below). This, in principle, hinders the robustness and generalization ability of deep learning models. Perhaps this is one of the reasons for which a general and robust model for habitat mapping in the Mediterranean Sea has been hitherto lacking. Thus, with this strict separation of training and out-of-sample test set we aim to comprehensively study the robustness and generalization power of our deep learning models.

Because we will finally train our model with all available data (from all regions), the images from 2023 will conform the final test set, to ensure the robustness of the model to changes in environmental conditions at the moment of image acquisition and image metadata.

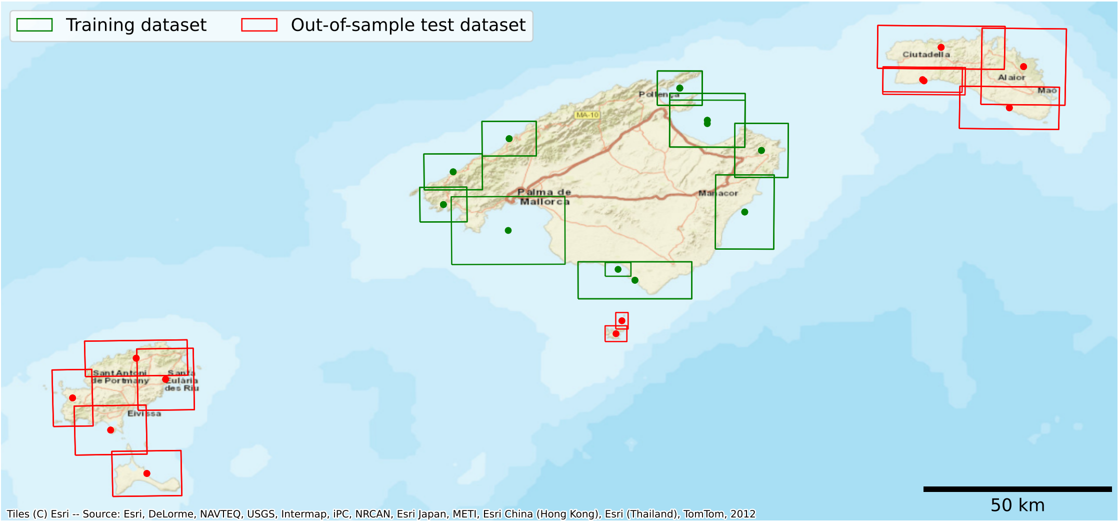


**Figure 1:** Coverage of the training and out-of-sample test datasets in the Balearic Islands. Each image was obtained for different years from 2020 to 2023, depending on the availability.

Table 1: Metadata of the satellite images used in the study.

| Name | Satellite azimuth | Sun azimuth | Sun elevation |
|---|---|---|---|
| Formentera_18_july_2021 | 99.4 | 113.5 | 57.3 |
| Formentera_24_july_2022 | 97.7 | 127.8 | 63 |
| Formentera_3_august_2023 | 110 | 128.8 | 60.5 |
| South_Oeste_Menorca_29_July_2022 | 100.6 | 116.1 | 54.1 |
| South_Oeste_Menorca_09_July_2021 | 182.4 | 114.2 | 58.1 |

Continued on next page

Table 1: Metadata of the satellite images used in the study. (Continued)

| | | | |
|---|---|---|---|
| Sur_Oeste_Menorca_9_august_2023 | 101.5 | 121.5 | 53.3 |
| Sur_Ibiza_20_july_2022 | 101.1 | 121.9 | 61.4 |
| Sur_Ibiza_18_july_2021 | 99.5 | 113.7 | 57.3 |
| Sur_Ibiza_26_august_2023 | 101.1 | 139.3 | 55.5 |
| Es_trenc_25_July_2022 | 270.7 | 113.8 | 54.6 |
| Es_Trenc_15_july_2023 | 112.8 | 112.6 | 56.7 |
| Norte_Menorca_29_July_2021 | 104.5 | 136.3 | 63 |
| Norte_Menorca_20_July_2022 | 176.1 | 118.6 | 58.1 |
| Norte_Menorca_23_june_2023 | 102 | 123.6 | 64.5 |
| Este_Menorca_19_July_2021 | 271.4 | 115.9 | 56.7 |
| Este_Menorca_15_July_2022 | 268 | 113.3 | 56.1 |
| Este_Menorca_23_june_2023 | 266.2 | 112.5 | 58.8 |
| CalaPi_CalaFiguera_29_july_2021 | 100.9 | 117.6 | 55.7 |
| CalaPi_CalaFiguera_23_july_2022 | 101.8 | 123.8 | 60.8 |
| CalaPi_CalaFiguera_12_july_2023 | 155.9 | 112.8 | 57.5 |
| PortoColom_CalaMillor_27_july_2021 | 278 | 117.4 | 55.9 |
| PortoColom_CalaMillor_22_july_2022 | 101.4 | 123.9 | 60.8 |
| PortoColom_CalaMillor_13_july_2023 | 101.7 | 113.1 | 57.5 |
| Sur_Este_menorca_02_July_2021 | 101.2 | 130.5 | 66.7 |
| Sur_Este_Menorca_30_July_2022 | 275.5 | 116.4 | 53.9 |
| Sur_Este_Menorca_9_august_2023 | 273.9 | 119.9 | 52.4 |
| Palma_7_july_2022 | 102.1 | 120.9 | 62.8 |
| Palma_29_june_2023 | 101.3 | 123.1 | 64.8 |
| Alcudia_25_July_2022 | 101 | 125.1 | 60.3 |
| Alcudia_21_May_2020 | 103.7 | 119.7 | 58.7 |
| Alcudia_22_July_2021 | 101.2 | 133.1 | 64.1 |
| Alcudia_15_july_2023 | 111.9 | 113.2 | 56.6 |
| Capdepera_20_july_2021 | 269.6 | 115.5 | 56.7 |
| Capdepera_22_july_2022 | 101.4 | 123.9 | 60.8 |
| Capdepera_31_july_2023 | 111.6 | 116.2 | 53.8 |
| Este_Ibiza_18_july_2021 | 99.6 | 113.8 | 57.3 |
| Este_Ibiza_24_july_2022 | 98.1 | 128.6 | 62.8 |
| Este_Ibiza_14_july_2023 | 277.5 | 124 | 63.3 |
| Pollença_6_July_2021 | 278.2 | 113.9 | 58.4 |
| Pollença_23_May_2020 | 277.1 | 119.7 | 58.8 |
| Pollença_21_July_2022 | 101.2 | 124.2 | 60.8 |
| Pollença_27_june_2023 | 106.9 | 123.5 | 64.4 |
| Oeste_Ibiza_18_july_2021 | 101.6 | 113.8 | 57.2 |
| Oeste_Ibiza_14_july_2022 | 101.3 | 120.8 | 62.3 |
| Oeste_Ibiza_19_july_2023 | 277.3 | 112.6 | 56.1 |

Continued on next page

Table 1: Metadata of the satellite images used in the study. (Continued)

| | | | |
|---|---|---|---|
| Banyalbufar_Soller_23_july_2021 | 176.3 | 116.8 | 56.5 |
| Banyalbufar_Soller_17_july_2022 | 110.5 | 123 | 61.5 |
| Banyalbufar_Soller_12_july_2023 | 101.5 | 124.5 | 63.2 |
| Dragonera_Banyalbufar_29_july_2022 | 102.8 | 120.4 | 56.9 |
| Dragonera_Banyalbufar_10_july_2021 | 110.5 | 113.7 | 58.1 |
| Dragonera_Banyalbufar_16_july_2023 | 101.1 | 125.8 | 63.1 |
| Norte_Ibiza_18_july_2021 | 101.6 | 113.8 | 57.2 |
| Norte_Ibiza_18_july_2022 | 277.9 | 112.7 | 56.2 |
| Norte_Ibiza_27_junio_2023 | 102 | 110.9 | 58.9 |
| South_Cabrera_23_july_2022 | 101.8 | 123.3 | 60.9 |
| South_Cabrera_29_june_2023 | 101.3 | 110.7 | 58.7 |
| North_Cabrera_19_july_2022 | 101.7 | 112 | 55.5 |
| North_Cabrera_14_july_2023 | 111.3 | 124 | 63.3 |

#### Ground truth dataset composition

The original seabed cartography contains a total of 28 different classes, which were aggregated into 4 major ecological groups or habitat types (*Posidonia oceanica*, Other green plants, Brown algae & rocks and Sandy bottoms) based on feature similarity and ecological function (Table 2). Although these habitat classes are present in the whole Mediterranean Sea, the specific composition of underlying sub-classes can vary among the different islands (e.g. one particular species of algae might be present in only one island, such as *Zostera noltii*).

**Table 2:** Ecological Categories and Subcategories in the ground truth habitat data.

| Category | Subcategory | Area ($km^2$) | Presence zone |
|---|---|---|---|
| **Posidonia oceanica** | Posidonia oceanica | 538.61 | Mallorca, Menorca, Ibiza, Formentera |
| | Barrier reef of Posidonia oceanica | 0.49 | Mallorca, Menorca |
| | Posidonia oceanica on stone with sand | 20.33 | Mallorca, Menorca |
| | Mixed Posidonia oceanica with dead rhizome | 0.10 | Ibiza |
| | Meadows of Posidonia oceanica on dead mat (rhizome) | 5.23 | Mallorca, Menorca |
| **Other Green Plants** | Algae photophilic on stone with Posidonia oceanica | 6.63 | Mallorca, Menorca |
| | Caulerpa prolifera | 0.82 | Mallorca, Menorca, Ibiza, Formentera |
| | Meadows of phanerogams and green rhizomatous algae | 4.87 | Mallorca, Menorca |
| | Fine sands with Cymodocea nodosa | 1.95 | Mallorca, Menorca, Ibiza, Formentera |
| | Cymodocea nodosa | 1.86 | Mallorca, Menorca, Ibiza, Formentera |
| | Zostera noltii | 0.01 | Menorca |
| | Cymodocea nodosa and Zostera noltii | 0.04 | Menorca |
| | Mixed meadows of Cymodocea nodosa and Caulerpa prolifera | 4.62 | Mallorca, Menorca, Ibiza |
| | Muddy bays with red algae (Alisidium corrallinum, Rytiphlaea tinctoria) | 0.01 | Menorca |
| **Sandy Bottoms** | Coarse sands | 250.19 | Ibiza, Formentera |
| | Soft or sedimentary substrate | 584.71 | Mallorca, Menorca |
| | Mud | 0.03 | Ibiza |
| | Fine sands | 74.36 | Ibiza, Formentera |
| | Muddy detrital bottom | 11.52 | Ibiza |
| | Leptometra phalangium fields in bathyal bottoms of platform edge | 208.85 | Menorca |
| | Bathyal bottoms of platform edge with Gryphus vitreus | 91.93 | Menorca |
| | Medium sands | 42.41 | Ibiza, Formentera |
| | Muddy detrital bottoms infralittoral and circalittoral | 589.81 | Mallorca, Menorca, Formentera |
| **Brown Algae and Rocks** | Rocky bottoms with photophilic algae and sands | 36.49 | Mallorca, Menorca |
| | Rocky bottoms dominated by sciafilic and hemisciafilic algae | 21.17 | Mallorca, Menorca, Ibiza, Formentera |
| | Cliffs, walls, and rocky slopes of the deep sea | 13.96 | Menorca |
| | Rocky bottoms with photophilic algae | 14.35 | Ibiza, Formentera |
| | Rocky bottoms with photophilic algae and sands | 15.58 | Mallorca, Menorca |

### Dataset creation

Satellite imagery, along with habitat and bathymetry data, were integrated to create comprehensive training and testing datasets for our model. Initially, the Near-Infrared (NIR) band was utilized to eliminate land pixels through a clustering algorithm, namely K-means, as this band is not able to penetrate water beyond 1 or 2 meters. Subsequently, the NIR band was replaced with bathymetry information. These processed satellite images served as the primary input data for our model. The ground truth dataset, or labels, consisted of raster files mirroring the satellite images, with single-band values indicating the benthic class for each pixel. Construction of this dataset involved associating each pixel in the processed satellite images with a corresponding benthic class based on aggregated habitat data. Pixels lacking a class assignment were masked out in both the satellite and label data. Similarly, pixels already masked in the satellite image were masked in the label image to ensure consistency.

Finally, patches of 256 x 256 pixels were created from each satellite and label image, forming the final dataset comprising up to 19369 patches. To study model performance in a real-case scenario, we trained our model with only data from the island of Mallorca (8488 patches, from which 1698 were used as validation set), while left as out-of-sample test set the data from the islands of Menorca, Ibiza, Formentera and Cabrera (7942 patches). We specifically call our test set "out-of-sample" test set to highlight the non-traditional way in which we test our model, which allow to test the extrapolation power and robustness of our model for real-case scenarios.

### Deep learning models

The performance of deep learning models can differ from one another due to its different architectures. Certain models can perform better than others in some situations, like segmenting specific classes or in images taken on different environmental conditions. Thus, we explored a selection of state-of-the-art deep learning models for semantic image segmentation such as UNET, Linknet, FPN and PSPNet (see below). Each model is formed by Convolutional Neural Network (CNN) blocks, designed to address specific challenges in semantic segmentation tasks. The specific architecture of the CNN blocks are usually referred to as the "backbone" of the model. We tested 10 different backbone models for each deep learning model, leading to the training and evaluation of 40 models.

#### UNET

UNET [1] is a popular architecture for image segmentation. It utilizes an encoder-decoder structure to capture high-level semantic information and detailed spatial features. The key feature of UNET is the inclusion of skip connections, enabling the model to propagate information from earlier layers to the corresponding decoder layers. This integration of skip connections allows the model to leverage both global context from the encoder and fine-grained spatial details from earlier layers, resulting in improved performance and accuracy. UNET's design aims to learn hierarchical representations while preserving spatial information, making it effective for various computer vision tasks.

#### Linknet

Linknet [2] follows an encoder-decoder structure and focuses on achieving a balance between accuracy and computational efficiency. Instead of traditional skip connections, Linknet incorporates "link blocks" to facilitate information flow between corresponding layers in the encoder and decoder paths. These link blocks consist of a shortcut connection and a residual connection, preserving and propagating important information during the upsampling process. Additionally, Linknet employs batch normalization and ReLU activation to enhance training convergence. The combination of skip connections, link blocks, and optimization techniques in Linknet results in improved information flow, enhanced spatial details, and reduced computational complexity.

#### FPN

FPN (Feature Pyramid Network) [3] is an architecture was introduced to tackle object detection and semantic segmentation across different scales. It addresses the challenge of capturing multi-scale information by constructing a feature pyramid with varying levels of feature maps. The architecture comprises a bottom-up pathway that extracts high-level semantic features using a CNN, such as ResNet or VGG, and a top-down pathway that generates feature maps by upsampling and merging information from higher-resolution levels. FPN incorporates lateral connections to combine low-level and high-level features, facilitating the fusion of fine-grained spatial details and high-level semantic information. The resulting feature pyramid enables effective detection and classification of objects of different sizes, making it well-suited multi-scale analysis classification problems.

#### PSPNet

PSPNet (Pyramid Scene Parsing Network) [4] is an architecture that utilizes a pyramid pooling module to capture contextual information at multiple scales. By dividing the input image into regions of varying sizes and aggregating global contextual information within each region, PSPNet improves the model's understanding of the scene and enhances object classification accuracy. The architecture comprises a convolutional neural network (CNN) backbone followed by the pyramid pooling module. This module performs pooling operations at different levels and spatial resolutions, capturing multi-scale context. The pooled features from each level are concatenated and passed to subsequent layers for classification. This enables PSPNet to effectively capture both local and global contextual information. The incorporation of the pyramid pooling mechanism enhances the model's comprehension of objects within the scene, ensuring robustness to variations in object scale and size.

#### Backbones

- **ResNet34**: A variant of ResNet with 34 layers, introducing residual learning to mitigate the vanishing gradient problem.
- **ResNet152**: A deeper variant of ResNet with 152 layers, capable of capturing more complex features.
- **SeResNet152 (SE-ResNet152)**: Based on ResNet, This architecture incorporates a Squeeze-and-Excitation block for adaptive recalibration of channel importance.
- **ResNeXt101**: While the ResNet model makes use of many smaller paths, ResNeXt substitutes "groups" for this function. There are several parallel pathways in these groupings, and distinct features are learned via each path.
- **SeResNeXt101 (SE-ResNeXt101)**: Combines ResNeXt architecture with Squeeze-and-Excitation for enhanced feature representation.
- **DenseNet201**: A Dense Convolutional Network connecting each layer in a feed-forward fashion for improved parameter efficiency.
- **InceptionV3**: Part of the Inception family, using parallel convolutional operations for features at various scales.
- **InceptionResNetV2**: Extends InceptionV3 with residual connections for improved training convergence.
- **EfficientNetB7**: Part of the EfficientNet family, balancing accuracy and efficiency for state-of-the-art performance using "Compound Scaling" methods.
- **MobileNetV2**: Optimized for mobile and edge devices, using depthwise separable convolutions for efficiency, to create deep neural networks that are lightweight and have minimal latency for embedded and mobile devices.

#### Performance Metrics

The evaluation of the deep learning models in our study involves the use of various performance metrics to assess their effectiveness in segmenting seagrass habitats. These metrics provide insights into the models' accuracy, precision, recall, F1 score, Cohen's kappa and intersection over union (IoU). Each metric serves a specific purpose in evaluating different aspects of model performance.

##### Accuracy

Accuracy measures the overall correctness of the model's predictions. It is calculated as the ratio of correctly predicted pixels to the total number of pixels in the dataset.

$$\text{Accuracy} = \frac{\text{True Positives} + \text{True Negatives}}{\text{Total Pixels}}$$

##### Precision

Precision is the ratio of true positive predictions to the total predicted positives, indicating how well the model performs when it predicts a certain class. Higher precision values imply fewer false positives.

$$\text{Precision} = \frac{\text{True Positives}}{\text{True Positives} + \text{False Positives}}$$

##### Recall

Recall, also known as sensitivity or true positive rate, measures the ability of the model to capture all instances of a given class. It is calculated as the ratio of true positive predictions to the total actual positives.

$$\text{Recall} = \frac{\text{True Positives}}{\text{True Positives} + \text{False Negatives}}$$

##### F1 Score

The F1 score is the harmonic mean of precision and recall. It provides a balanced measure of a model's performance, especially when dealing with imbalanced datasets. A higher F1 score indicates better overall performance.

$$\text{F1 Score} = 2 \times \frac{\text{Precision} \times \text{Recall}}{\text{Precision} + \text{Recall}} = \frac{2 \times \text{True Positives}}{2 \times \text{True Positives} + \text{False Negatives} + \text{False Positives}}$$

##### Cohen's Kappa

Cohen's Kappa is a statistical measure of inter-rater agreement for categorical items. It is generally thought to be a more robust measure than simple percent agreement calculation, as Kappa takes into account the possibility of the agreement occurring by chance.

$$\kappa = \frac{p_o - p_e}{1 - p_e}$$

where $p_o$ is the relative observed agreement among raters, and $p_e$ is the hypothetical probability of chance agreement, using the observed data to calculate the probabilities of each observer randomly seeing each category.

##### Intersection over Union (IoU)

IoU, also known as Jaccard Index, measures the overlap between the predicted and true positive pixels. It is calculated as the ratio of the intersection of predicted (P) and label (L) pixels to the union of all pixels. IoU is particularly useful for semantic segmentation tasks, providing insights into the spatial accuracy of predictions.

$$\text{IoU} = \frac{|P \cap L|}{|P \cup L|} = \frac{|P \cap L|}{|P| + |L| - |P \cap L|}$$

#### Spectral reflectance analysis

To gain a little understanding on our problem, we first perform a basic analysis on the spectral reflectance of the habitat classes. The distribution of reflectance values among the spectral bands plays a crucial role in the ability of the model to segment the different classes. When the reflectance values of the different classes do not overlap, the discrimination capability of the machine learning model is enhanced. Conversely, when reflectance values show significant overlap between classes, the model may encounter difficulties in categorizing pixels accurately, as the spectral information becomes less informative.

We computed the distribution of the response values of each habitat class to each of the satellite bands together with the mean Wasserstein distance among the distributions (Fig. 2). The Blue, Green and Green 1 bands are highlighted as potentially more informative for the model, followed by Coastal Blue and Yellow. Red, Red Edge and NIR are identified as the bands with potentially less discrimination power. This could be expected from the fact that the wavelengths at the extreme ends of the visible spectrum are attenuated faster than those wavelengths in the middle. Finally, we observe that depth data is, by itself, basically uninformative.

Of course, this is just a simple, rather linear, analysis of the information provided to the model by the satellite images. In the end, the AI models will try to segment the different classes based on the representation of the response values in an abstract hyperspace.

Something interesting is that depth has a very low discrimination power alone, but, we advance, observed that introducing the depth to the model highly improves model predictions. Thus, we can hypothesize that the model is somehow learning and applying a kind of depth reflectance correction to the input data to increase its accuracy.

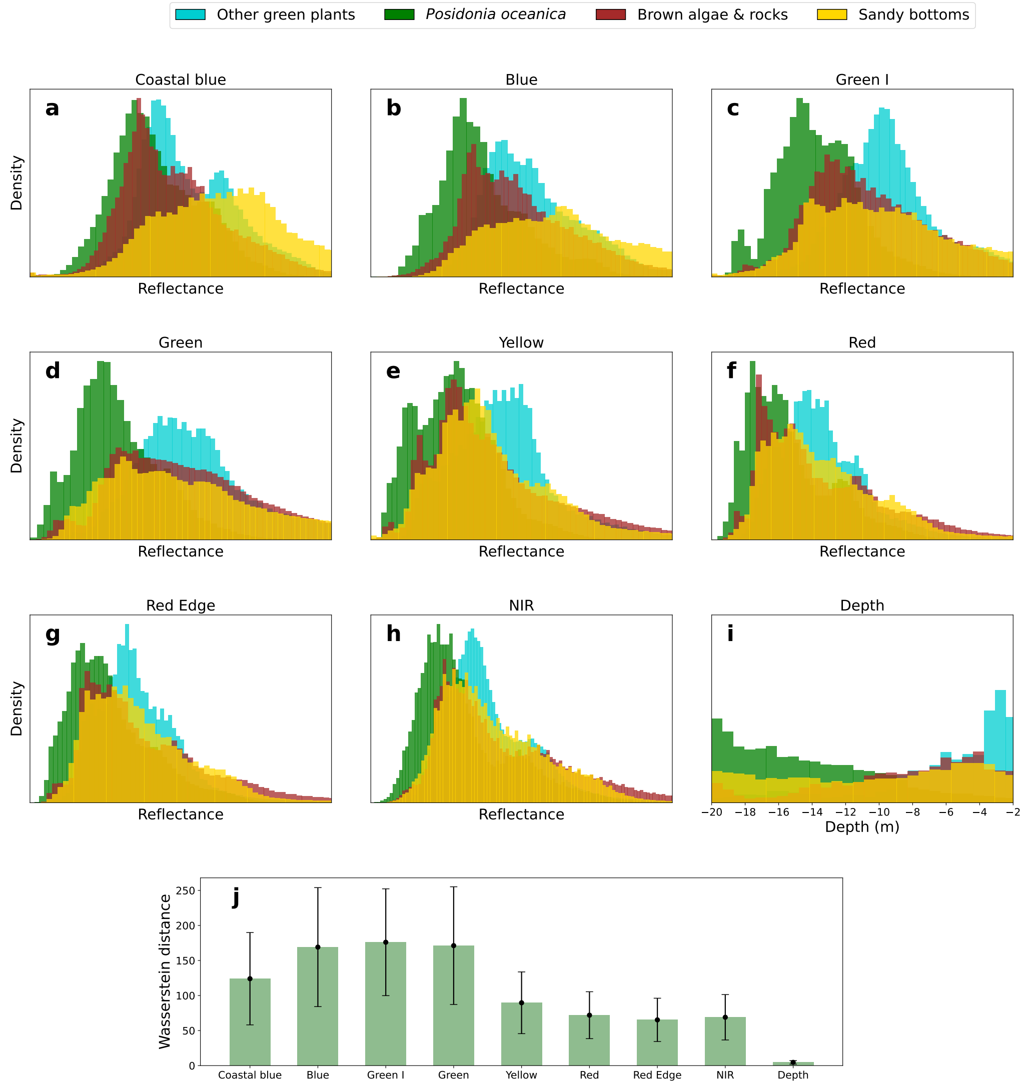


**Figure 2:** Distribution of response values (surface reflectance) for the different processed habitat classes with respect to the different bands available from the satellite imagery and depth.

### Architecture selection

We trained the 40 deep learning models as specified in the Methods section. After training, the performance on both train and validation sets was compared among the models. Models based on UNET and Linknet clearly outperformed models based on PSPNet and FPN architectures (Table 3). Despite the similar performance of the models based on both UNET and Linknet architectures, UNET is a more complex architecture than Linknet, which translates into a bigger number of trainable parameters (Table 4). Thus, models based on UNET architecture are more computationally expensive and, in addition, are more prone to suffer from over-fitting. Thus, we finally selected Linknet as the main architecture for our models.

**Table 3:** Performance among all architectures for all backbones based on the Intersection over Union metric.

| Architecture | UNET | Linknet | PSPNet | FPN |
|---|---|---|---|---|
| densenet201 | 91.15 | **91.51** | 86.93 | 87.97 |
| resnet152 | **92.01** | 91.95 | 86.94 | 90.64 |
| seresnext101 | 91.52 | **91.54** | 85.52 | 87.88 |
| efficientnetb7 | **89.27** | 88.65 | 85.99 | 87.19 |
| inceptionv3 | 89.49 | **90.59** | 85.13 | 89.33 |
| seresnet152 | 91.76 | **91.83** | 86.49 | 90.48 |
| inceptionresnetv2 | 91.85 | **91.91** | 86.16 | 89.61 |
| resnext101 | 92.16 | **92.5** | 86.96 | 89.09 |
| mobilenetv2 | **89.35** | 89.16 | 83.15 | 89.67 |
| resnet34 | **90.44** | 90.14 | 86.12 | 90.05 |

**Table 4:** Number of total parameters among all architectures for all backbones (in millions)

| Architecture | UNET | Linknet |
|---|---|---|
| resnet152 | 67.31 | **63.53** |
| mobilenetv2 | 8.04 | **4.14** |
| seresnet152 | 73.95 | **70.17** |
| inceptionv3 | 29.93 | **26.27** |
| densenet201 | 26.39 | **22.56** |
| inceptionresnetv2 | 62.06 | **57.87** |
| seresnext101 | 56.07 | **52.29** |
| efficientnetb7 | 75.05 | **72.26** |
| resnet34 | 24.47 | **21.65** |
| resnext101 | 51.29 | **47.52** |

### Out-of-sample predicting power and robustness

Because the performance of the different backbones can differ from one another, with some being better than others in certain situations, we implemented a pixel-wise consensus algorithm to enhance the robustness and reliability of model predictions. Basically, the results from all models were aggregated and the class label with the highest frequency across all predictions was assigned to the each pixel. Following this algorithm, we tested different aggregation strategies: selecting all the models (voting_all), selecting only the best 3 or 5 models (voting_top_3, voting_top_5) and selecting the models that performed better in segmenting each class individually (voting_specialists).

**Table 5:** Performance metrics for all models based on Linknet architecture in the training dataset

| Backbone | IoU | f1 | Kappa | Precision | Recall | Accuracy |
|---|---|---|---|---|---|---|
| **voting_top_3** | **89.57** | **94.01** | **80.72** | **94.79** | **94.37** | **94.28** |
| voting_top_5 | 89.01 | 93.64 | 80.12 | 94.59 | 94.01 | 93.9 |
| inceptionresnetv2 | 88.95 | 93.72 | 81.35 | 94.3 | 93.99 | 93.9 |
| resnext101 | 88.46 | 93.27 | 78.48 | 94.22 | 93.65 | 93.55 |
| resnet152 | 88.34 | 93.23 | 78.72 | 94.12 | 93.62 | 93.54 |
| voting_all | 88.22 | 93.13 | 78.63 | 94.24 | 93.56 | 93.44 |
| voting_specialists | 88.22 | 93.21 | 79.27 | 94.2 | 93.57 | 93.46 |
| mobilenetv2 | 88.01 | 93.16 | 80.74 | 93.78 | 93.37 | 93.28 |
| resnet34 | 86.43 | 91.95 | 75.29 | 93.12 | 92.47 | 92.33 |
| inceptionv3 | 85.96 | 91.74 | 77.56 | 92.88 | 92.09 | 92.04 |
| seresnet152 | 85.74 | 91.54 | 76.36 | 92.94 | 91.91 | 91.79 |
| seresnext101 | 85.65 | 91.55 | 76.2 | 92.75 | 91.92 | 91.77 |
| efficientnetb7 | 84.56 | 90.79 | 72.95 | 91.86 | 91.35 | 91.19 |
| densenet201 | 82.21 | 89.04 | 69.72 | 90.99 | 89.67 | 89.48 |
| Median | 88.12 | 93.16 | 78.56 | 93.95 | 93.47 | 93.36 |

**Table 6:** Performance metrics for all models based on Linknet architecture in the out-of-sample test dataset

| Backbone | IoU | f1 | Kappa | Precision | Recall | Accuracy |
|---|---|---|---|---|---|---|
| **efficientnetb7** | **62.19** | **73.05** | **54.34** | 76.1 | 74.78 | 74.62 |
| **voting_all** | 61.97 | 72.77 | 53.7 | **76.74** | **75.02** | **74.88** |
| inceptionresnetv2 | 61.80 | 72.69 | 52.8 | 75.31 | 74.87 | 74.7 |
| voting_top_5 | 61.35 | 72.38 | 53.11 | 76.29 | 74.38 | 74.25 |
| voting_specialists | 61.10 | 71.92 | 52.2 | 75.93 | 74.47 | 74.37 |
| voting_top_3 | 60.88 | 71.92 | 52.03 | 75.94 | 74.08 | 73.94 |
| mobilenetv2 | 60.87 | 72.01 | 51.74 | 74.40 | 74.12 | 73.97 |
| seresnext101 | 60.58 | 71.82 | 52.02 | 75.01 | 73.57 | 73.35 |
| resnext101 | 60.46 | 71.68 | 51.64 | 75.39 | 73.68 | 73.50 |
| seresnet152 | 60.14 | 71.48 | 52.77 | 75.58 | 72.94 | 72.67 |
| inceptionv3 | 59.65 | 70.82 | 50.66 | 73.81 | 72.85 | 72.65 |
| densenet201 | 59.2 | 70.69 | 51.57 | 75.37 | 72.01 | 71.83 |
| resnet152 | 57.58 | 69.16 | 48.89 | 73.04 | 71.00 | 70.91 |
| resnet34 | 57.54 | 69.2 | 47.38 | 72.54 | 71.32 | 71.16 |
| Median | 60.73 | 71.87 | 52.03 | 75.38 | 73.88 | 73.72 |

We compared the performance of our Linknet models and consensus strategies in both the training and out-of-sample test datasets. Regarding the training set, we observe that some models perform better than others, as expected. At an individual basis, inceptionresnetv2 and resnext101 are the best-performing backbones, while efficientnetb7 and densenet201 are the worst-performing ones (Table 5). The consensus strategies cluster at the first positions of the ranking, with voting_top_3 and voting_top_5 occupying the two first positions. Interestingly, we observe that efficientnetb7 is the best-performing backbone in the test set, followed by voting_all and inceptionresnetv2 (Table 6). This suggests that efficientnetb7 is one of the most robust models, with higher generalization power. At any rate, we observe that it is closely followed by the voting_all consensus strategy. Here we must emphasize that while the mask class is considered during training, it is omitted during evaluation and prediction phases to focus solely on the specified classes of interest. Consequently, this contributes to the observed differences in IoU values between Table 3 and Tables 5 and 6.

Overall, we note that the relative performance of the model in the out-of-sample test set is significantly reduced in comparison with the training set, although the absolute performance is still considerable. This, indeed, is expected, as the habitat classes of the out-of-sample test are not formed by exactly the same sub-categories that in the training set. Furthermore, there is a huge environmental variability and intrinsic noise in the overall framework proposed here: the relative position of the satellite with respect to the sun and the earth at the time of each image acquisition, the specific atmospheric and marine conditions, the fact that some species forming the habitat classes are seasonal, etc.

#### Understanding model performance

**Table 7:** Performance in segmenting each habitat class individually, measured by Intersection over Union, for all models in training and out-of-sample test datasets.

| | Training dataset | | | | Test dataset | | | |
|---|---|---|---|---|---|---|---|---|
| Method | PO | OGP | SB | BAR | PO | OGP | SB | BAR |
| densenet201 | 88.69 | 69.9 | 75.35 | 66.25 | 72.48 | 5.44 | 55.12 | 38.24 |
| efficientnetb7 | 90.26 | 69.56 | 77.37 | 74.66 | 76.99 | 3.86 | **56.31** | 40.59 |
| inceptionresnetv2 | **92.61** | **85.81** | 83.63 | 81.17 | **77.34** | 3.72 | 53.85 | **41.17** |
| inceptionv3 | 90.8 | 73.6 | 80.82 | 75.57 | 75.79 | 3.79 | 50.99 | 38.21 |
| mobilenetv2 | 92.05 | 83.6 | 81.49 | 81.44 | 76.76 | **5.49** | 53.16 | 37.27 |
| resnet152 | 91.87 | 85.61 | 82.79 | **81.7** | 71.54 | 2.92 | 51.04 | 37.92 |
| resnet34 | 90.8 | 82.7 | 80.78 | 75.84 | 73.11 | 3.37 | 49.71 | 35.7 |
| resnext101 | 92.09 | 85.78 | **83.65** | 79.56 | 75.74 | 4.39 | 52.92 | 39.6 |
| seresnet152 | 90.87 | 75.98 | 79.63 | 74.45 | 73.09 | 4.82 | 55.65 | 40.4 |
| seresnext101 | 90.67 | 77.45 | 78.53 | 76.53 | 75.61 | 4.09 | 53.69 | 39.49 |
| voting_all | 91.97 | 85.29 | 82.2 | 81.54 | 77.30 | **4.24** | **54.50** | **41.99** |
| voting_top_3 | **92.79** | **87.28** | **84.72** | **83.01** | 76.25 | 4.12 | 53.11 | 40.57 |
| voting_top_5 | 92.5 | 86.1 | 83.54 | 82.53 | 76.18 | 4.13 | 54.44 | 41.71 |
| voting_specialists | 92.01 | 86.78 | 82.38 | 80.21 | **77.31** | 4.22 | 51.88 | 40.69 |
| Area in ($km^2$) | 359.96 | 29.31 | 139.46 | 63.82 | 156.12 | 18.17 | 80.59 | 36.96 |

We then studied the performance of the model on segmenting each of the ecological habitats individually. We observed the emergence of “specialists”: in the training set, inceptionresnet outperforms in segmenting *Posidonia oceanica* meadows and Other green plants, resnext101 is better suited for mapping Sandy bottoms while resnet152 shows enhanced performance for detecting Brown algae & rocks (Table 7).

However, the results change when looking at the test dataset. This somehow explains why the voting_all consensus method outperforms the voting_smart one. We also note that the performance of the model with respect to Other green plants is drastically reduced in the test set, which undoubtedly affect the overall performance of the model shown in Table 6.

To further understand this drastic loss of performance on segmenting the "Other green plants" class, we computed the confusion matrix for the model (voting_all from now on) predictions (Fig. 3). In the training dataset there is already a significant difference between the performance in segmenting *Posidonia oceanica* meadows, with 99.5% True Positives (TP) and the other classes, with 89.4% TP for Other green plants, 84.5% TP for Brown algae & rocks and 83.6% TP for Sandy Bottoms (Fig. 3 a). At any rate, those differences clearly increase in the out-of-sample test set. While the segmentation of *Posidonia oceanica* meadows keeps a remarkable 94.6% TP, with a still low FP rate, the TP rate for the other classes is reduced (Fig. 3 b). Specifically, a significant part of the pixels that were categorized by the ground truth data as Other green plants, Brown algae & rocks or Sandy bottoms are being classified by the model as *Posidonia oceanica* meadows. This is the reason for the very low IoU score of the Other green plant class in the testing set and the overall IoU decrease in all classes. In addition, we can also observe that the area of each class is significantly reduced in the test set in comparison with the training set (Fig. 3c-d, sum over rows).

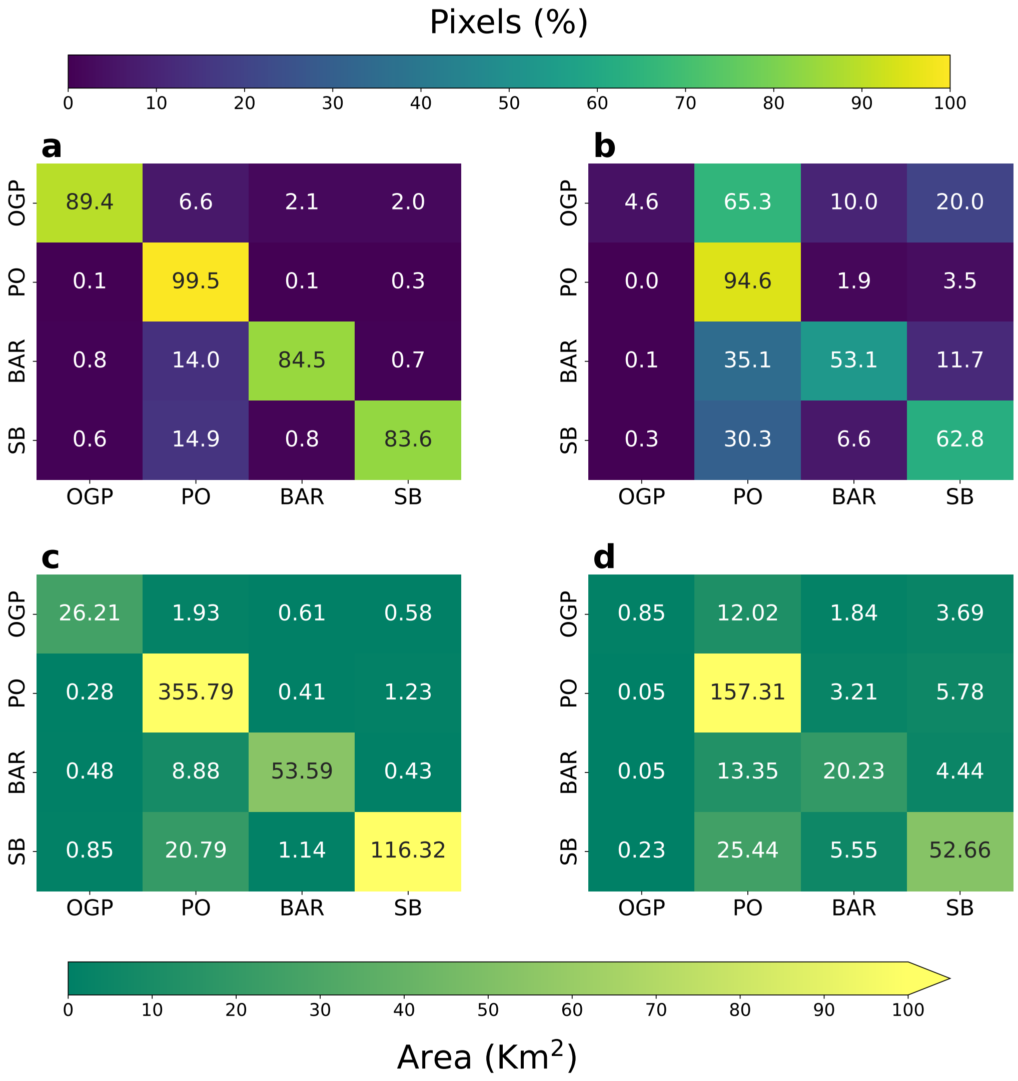


**Figure 3:** Confusion matrix for train (a) and out-of-sample test (b) dataset. Confusion matrix for the predicted area in train (c) and out-of-sample test (d) datasets.

To better understand this behavior, we compared the response values of each class between train and test datasets. In principle, the distribution of response values for a given class in the test set should be similar to the distribution of that class in the training set. This is indeed the rationale behind the use of correlative models. Of course, this distributions should be similar in a high dimensional space that we are not able to visualize. At any rate, to gain some understanding, we computed the wasserstein distance between the distribution of response values in train and test for the Blue, Green and Green I bands, i.e. the most influential bands, as previously found.

The results revealed that the distribution of response values for the Other green plants class in the test dataset is much more similar to the *Posidonia oceanica* distribution of response values in the train set than to its own class (Fig. 4), specially for Green I and Green bands (Fig. 4 b-c). Then it is not surprising that the model classifies all those samples as *Posidonia oceanica*. Indeed, because of the great seasonality and natural variability of some of the species conforming the Other green plants class, the real habitat class present at the time of the satellite image acquisition could deviate from the initial one present at sonar-based classification. Similarly, there are some mixed sub-classes such as "Photophilic algae on stone with *Posidonia oceanica*" which further increase the probability of the previous argument taking place. That would explain the fact that the distribution of the response values for the "Other green plants" class in the test dataset is more similar to the *Posidonia oceanica* distribution in the training set than to its own class in the training set. So, in summary, it is plausible that some of the model predictions that are supposed to be incorrect, were indeed correct.

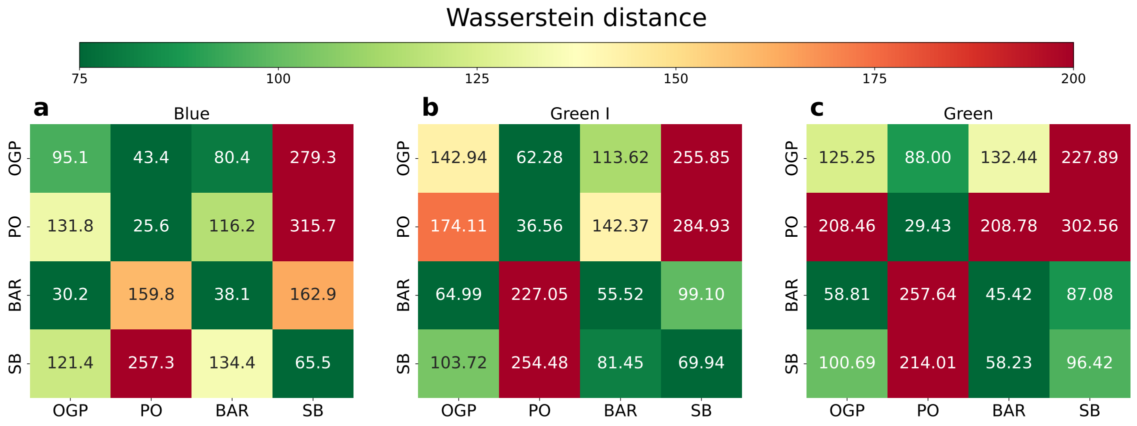


**Figure 4:** Wasserstein distance between Train and Test datasets for Blue (a), Green I (b) and Green (c) bands.

All the analysis on model performance done up to this point were based on **mean** results over the training and out-of-sample test datasets, which are based on 21 images each. Now we analyze the performance of the model predictions in each image individually (Table 8). We observe that there is a great variability in model performance within each dataset. In the training set, 60% of the images are segmented with IoU> 90% and 30% with IoU> 80%. The resting 2 images, however, are segmented with "only" around 64% IoU score, overlapping with the out-of-sample test dataset (Table 8). In the test set, 15% of the images are segmented with IoU> 80%, 20% with IoU> 70% and 45% with IoU> 50%. The resting 4 images are segmented with an IoU score of around 40%. In any case, the accuracy of all segmentations are always over 50%, much above the threshold for random segmentation of 25% (recall it is a 4-class classification problem).

These results present two interesting insights. First, the model is able to segment some images in the test set with very high performance (15% of the images). Recall this is not a usual testing set, but we have only trained the model in a given island (Mallorca) and tested it in 3 other islands (Menorca, Ibiza and Formentera), not controlling for neither weather or satellite-position variability. Second, we observe that 2 images in the training test are segmented with rather "low" performance, situating at the middle of the testing set in the performance ranking (Table 8). Interestingly, those are images situated in the "Serra de Tramuntana", the mountain range in Mallorca. The sea bottom in this area is characterized by a rocky substrate where depth increases rapidly. Indeed, we can observe that the images that are segmented with higher performance in the test set (Formentera_24_july_2022 to Sur_Ibiza_20_july_2022) are precisely characterized by more sandy and clear sea bottoms with slowly increasing depths, while the rest are rather rocky with rapidly increasing depth (Table 8). This insights can help in taking future steps to develop a general model for segmenting sea habitats in the whole Mediterranean sea.

**Table 8:** Performance metrics for voting_all model in segmenting individual images from training and testing datasets.

| **Image Name** | **IoU** | **f1** | **Kappa** | **Precision** | **Recall** | **Accuracy** | **Area ($Km^2$)** | **Dataset** |
|---|---|---|---|---|---|---|---|---|
| PortoColom_CalaMillor_22_july_2022 | 94.77 | 97.27 | 94.21 | 97.31 | 97.3 | 97.24 | 28.22 | Train |
| Es_trenc_25_july_2022 | 93.56 | 96.53 | 87.1 | 96.81 | 96.57 | 96.54 | 12.65 | Train |
| Capdepera_22_july_2022 | 93.23 | 96.43 | 94.04 | 96.52 | 96.47 | 96.37 | 15.89 | Train |
| Pollença_6_july_2021 | 92.78 | 96.06 | 86.42 | 96.3 | 96.27 | 96.22 | 26.02 | Train |
| CalaPi_CalaFiguera_23_july_2022 | 92.34 | 95.83 | 90.89 | 96.12 | 96.02 | 95.96 | 36.58 | Train |
| Alcudia_25_july_2022 | 91.97 | 95.52 | 79.92 | 95.92 | 95.83 | 95.81 | 76.18 | Train |
| Capdepera_20_july_2021 | 91.37 | 95.36 | 92.31 | 95.6 | 95.43 | 95.34 | 15.89 | Train |
| Pollença_23_may_2020 | 91.12 | 94.99 | 81.0 | 95.44 | 95.39 | 95.33 | 24.7 | Train |
| CalaPi_CalaFiguera_29_july_2021 | 90.87 | 94.95 | 90.16 | 95.47 | 95.15 | 95.09 | 36.53 | Train |
| Banyalbufar_Soller_23_july_2021 | 90.38 | 94.89 | 90.71 | 95.07 | 94.94 | 94.81 | 8.88 | Train |
| Dragonera_Toro_23_july_2021 | 90.06 | 94.63 | 90.71 | 94.82 | 94.76 | 94.5 | 11.93 | Train |
| Dragonera_Toro_22_july_2022 | 90.05 | 94.65 | 90.62 | 94.84 | 94.77 | 94.49 | 11.87 | Train |
| Pollença_21_july_2022 | 89.82 | 94.01 | 77.29 | 94.83 | 94.7 | 94.65 | 26.05 | Train |
| PortoColom_CalaMillor_27_july_2021 | 88.91 | 93.99 | 86.49 | 94.44 | 94.14 | 94.08 | 28.22 | Train |
| Alcudia_22_july_2021 | 86.06 | 91.52 | 63.17 | 93.14 | 92.51 | 92.47 | 75.69 | Train |
| Alcudia_21_may_2020 | 84.62 | 90.44 | 58.93 | 92.69 | 91.34 | 91.3 | 73.63 | Train |
| Dragonera_Banyalbufar_10_july_2021 | 83.97 | 91.04 | 85.27 | 92.17 | 91.11 | 90.87 | 7.09 | Train |
| Formentera_24_july_2022 | 82.57 | 89.53 | 82.24 | 89.73 | 89.74 | 88.55 | 28.96 | Test |
| Formentera_18_july_2021 | 82.54 | 89.52 | 82.27 | 89.9 | 89.84 | 88.68 | 28.99 | Test |
| Palma_7_july_2022 | 82.42 | 90.03 | 81.53 | 91.59 | 90.1 | 88.99 | 57.4 | Train |
| Sur_Oeste_Menorca_09_July_2021 | 82.12 | 88.25 | 58.03 | 87.84 | 89.5 | 89.55 | 10.87 | Test |
| North_Cabrera_19_july_2022 | 77.78 | 86.24 | 49.14 | 87.42 | 85.54 | 85.54 | 1.37 | Test |
| Sur_Oeste_Menorca_29_July_2022 | 75.14 | 84.15 | 36.85 | 88.46 | 81.84 | 80.29 | 16.4 | Test |
| Sur_Ibiza_18_july_2021 | 70.41 | 79.92 | 51.71 | 80.02 | 81.73 | 82.14 | 21.47 | Test |
| Sur_Ibiza_20_july_2022 | 70.11 | 79.64 | 50.29 | 79.84 | 81.55 | 81.97 | 22.52 | Test |
| South_Cabrera_23_july_2022 | 64.89 | 77.3 | 47.39 | 81.76 | 76.02 | 76.0 | 1.45 | Test |
| Banyalbufar_Soller_17_july_2022 | 64.49 | 76.47 | 54.73 | 84.17 | 79.5 | 79.4 | 8.83 | Train |
| Dragonera_Banyalbufar_29_july_2022 | 63.14 | 76.1 | 62.49 | 85.53 | 76.27 | 75.88 | 7.25 | Train |
| Este_Ibiza_18_july_2021 | 62.28 | 73.4 | 43.37 | 77.75 | 76.52 | 76.53 | 13.19 | Test |
| Este_Ibiza_24_july_2022 | 61.53 | 72.59 | 42.15 | 77.8 | 75.62 | 75.67 | 13.21 | Test |
| Oeste_Ibiza_14_july_2022 | 59.56 | 73.26 | 54.19 | 76.74 | 75.17 | 75.92 | 7.72 | Test |
| Oeste_Ibiza_18_july_2021 | 58.9 | 72.63 | 53.26 | 76.26 | 74.69 | 75.59 | 7.75 | Test |
| Sur_Este_Menorca_29_July_2022 | 58.0 | 69.34 | 57.21 | 67.93 | 74.34 | 75.16 | 16.15 | Test |
| Sur_Este_Menorca_02_July_2021 | 57.01 | 68.53 | 55.82 | 71.64 | 73.55 | 74.38 | 16.42 | Test |
| Norte_Ibiza_18_july_2022 | 53.01 | 68.57 | 53.45 | 70.57 | 69.52 | 69.21 | 4.27 | Test |
| Norte_Ibiza_18_july_2021 | 50.03 | 66.05 | 49.13 | 71.43 | 66.85 | 67.13 | 4.4 | Test |
| Norte_Menorca_20_July_2022 | 43.32 | 55.8 | 43.98 | 69.5 | 58.77 | 58.33 | 13.85 | Test |
| Este_Menorca_19_July_2021 | 42.26 | 56.51 | 42.5 | 61.63 | 60.13 | 60.21 | 25.49 | Test |
| Este_Menorca_15_July_2022 | 38.35 | 52.87 | 36.78 | 63.33 | 57.53 | 57.42 | 25.95 | Test |
| Norte_Menorca_29_July_2021 | 38.28 | 51.3 | 37.52 | 67.92 | 54.69 | 54.37 | 14.23 | Test |

#### Effect of depth on model performance

Finally, we studied the effect of depth in model performance, focusing on the test set. We observe that, in general, model performance is rather independent on depth (Fig. 5 a). Interestingly, we observe that the *Posidonia oceanica* class consistently exhibits the highest IoU values across all depths compared to other classes. This noteworthy trend may be attributed to the abundance of *Posidonia oceanica* samples (Fig. 5 b), suggesting that the model's performance excels in accurately identifying and delineating Posidonia in underwater environments.

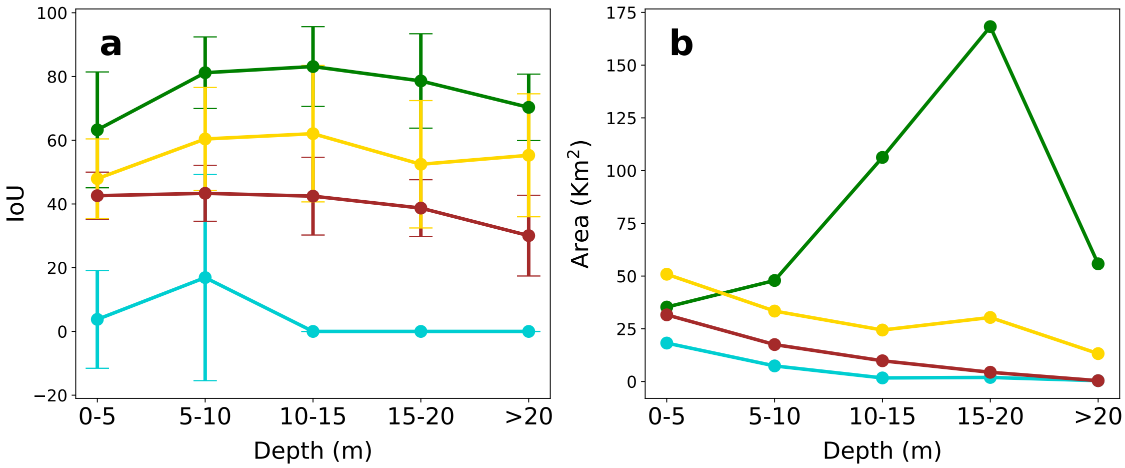


**Figure 5:** (a) Performance of voting_all model as function of the depth of classified pixels. (b) Area of samples contained in each depth range for the different habitat classes.

### CAMELE trained with all available data

**Table 9:** Performance metrics for voting_all model in segmenting individual images from previous training and testing datasets.

| Image Name | IoU | f1 | Kappa | Precision | Recall | Accuracy | Area ($Km^2$) | Dataset |
|---|---|---|---|---|---|---|---|---|
| Formentera_18_july_2021 | 98.50 | 99.24 | 98.50 | 99.25 | 99.25 | 99.25 | 29.07 | Test |
| Formentera_24_july_2022 | 98.42 | 99.20 | 98.43 | 99.20 | 99.20 | 99.20 | 29.04 | Test |
| Sur_Oeste_Menorca_09_July_2021 | 97.35 | 98.62 | 93.39 | 98.65 | 98.65 | 98.65 | 11.01 | Test |
| Sur_Oeste_Menorca_29_July_2022 | 97.33 | 98.59 | 91.86 | 98.64 | 98.65 | 98.65 | 16.40 | Test |
| Sur_Ibiza_20_july_2022 | 96.74 | 98.32 | 95.88 | 98.34 | 98.34 | 98.34 | 22.60 | Test |
| Es_trenc_25_july_2022 | 96.70 | 98.28 | 92.34 | 98.31 | 98.31 | 98.31 | 12.65 | Train |
| Sur_Ibiza_18_july_2021 | 96.69 | 98.29 | 95.99 | 98.31 | 98.31 | 98.31 | 21.67 | Test |
| Norte_Menorca_20_July_2022 | 96.46 | 98.19 | 97.36 | 98.20 | 98.20 | 98.20 | 14.13 | Test |
| Este_Menorca_19_July_2021 | 96.04 | 97.97 | 97.05 | 97.99 | 97.98 | 97.98 | 27.18 | Test |
| CalaPi_CalaFiguera_29_july_2021 | 96.03 | 97.94 | 95.71 | 97.97 | 97.97 | 97.97 | 36.63 | Train |
| Este_Menorca_15_July_2022 | 96.03 | 97.97 | 97.04 | 97.99 | 97.97 | 97.97 | 27.04 | Test |
| CalaPi_CalaFiguera_23_july_2022 | 96.01 | 97.94 | 95.70 | 97.96 | 97.96 | 97.96 | 36.59 | Train |
| PortoColom_CalaMillor_22_july_2022 | 95.85 | 97.85 | 95.50 | 97.87 | 97.87 | 97.87 | 28.22 | Train |
| Sur_Este_Menorca_29_July_2022 | 95.80 | 97.84 | 96.59 | 97.85 | 97.85 | 97.85 | 16.54 | Test |
| Sur_Este_Menorca_02_July_2021 | 95.79 | 97.83 | 96.58 | 97.85 | 97.84 | 97.84 | 16.52 | Test |
| Norte_Menorca_29_July_2021 | 95.69 | 97.79 | 96.76 | 97.81 | 97.79 | 97.79 | 14.25 | Test |
| Palma_7_july_2022 | 95.68 | 97.78 | 95.95 | 97.79 | 97.78 | 97.78 | 57.39 | Train |
| PortoColom_CalaMillor_27_july_2021 | 95.55 | 97.69 | 95.15 | 97.71 | 97.71 | 97.71 | 28.22 | Train |
| Alcudia_25_july_2022 | 95.26 | 97.47 | 89.41 | 97.57 | 97.57 | 97.57 | 76.19 | Train |
| North_Cabrera_19_july_2022 | 95.25 | 97.48 | 89.80 | 97.50 | 97.53 | 97.53 | 1.38 | Test |
| Alcudia_21_may_2020 | 94.88 | 97.23 | 87.03 | 97.35 | 97.37 | 97.37 | 73.63 | Train |
| Alcudia_22_july_2021 | 94.80 | 97.21 | 88.23 | 97.34 | 97.33 | 97.33 | 75.70 | Train |
| Capdepera_22_july_2022 | 94.64 | 97.19 | 95.32 | 97.22 | 97.22 | 97.22 | 15.90 | Train |
| Capdepera_20_july_2021 | 94.58 | 97.15 | 95.28 | 97.19 | 97.19 | 97.19 | 15.92 | Train |
| Este_Ibiza_24_july_2022 | 94.02 | 96.88 | 92.80 | 96.93 | 96.91 | 96.91 | 13.28 | Test |
| Pollença_23_may_2020 | 94.01 | 96.80 | 89.11 | 96.89 | 96.91 | 96.91 | 24.73 | Train |
| Este_Ibiza_18_july_2021 | 93.85 | 96.79 | 92.54 | 96.83 | 96.82 | 96.82 | 13.29 | Test |
| Pollença_21_july_2022 | 93.41 | 96.46 | 88.34 | 96.59 | 96.59 | 96.59 | 26.05 | Train |
| Oeste_Ibiza_14_july_2022 | 93.34 | 96.51 | 94.01 | 96.57 | 96.55 | 96.55 | 7.76 | Test |
| Oeste_Ibiza_18_july_2021 | 93.34 | 96.51 | 94.00 | 96.57 | 96.56 | 96.56 | 7.77 | Test |
| Pollença_6_july_2021 | 92.71 | 96.07 | 87.63 | 96.19 | 96.21 | 96.21 | 26.05 | Train |
| Banyalbufar_Soller_23_july_2021 | 92.64 | 96.14 | 93.06 | 96.20 | 96.17 | 96.17 | 8.89 | Train |
| Banyalbufar_Soller_17_july_2022 | 92.61 | 96.12 | 93.02 | 96.19 | 96.15 | 96.15 | 8.88 | Train |
| Dragonera_Banyalbufar_29_july_2022 | 92.42 | 95.91 | 93.47 | 96.04 | 96.04 | 96.04 | 7.28 | Train |
| Dragonera_Banyalbufar_10_july_2021 | 92.35 | 95.88 | 93.41 | 96.01 | 96.00 | 96.00 | 7.25 | Train |
| Norte_Ibiza_18_july_2022 | 92.04 | 95.74 | 93.75 | 95.91 | 95.85 | 95.85 | 4.40 | Test |
| South_Cabrera_23_july_2022 | 91.75 | 95.56 | 89.64 | 95.52 | 95.66 | 95.66 | 1.53 | Test |
| Norte_Ibiza_18_july_2021 | 90.76 | 95.01 | 92.67 | 95.26 | 95.13 | 95.13 | 4.41 | Test |
| Dragonera_Toro_23_july_2021 | 90.48 | 94.88 | 91.19 | 95.05 | 94.99 | 94.99 | 11.93 | Train |
| Dragonera_Toro_22_july_2022 | 89.95 | 94.58 | 90.50 | 94.78 | 94.71 | 94.71 | 11.93 | Train |

**Table 10:** Performance metrics for the final model in segmenting individual images for the year 2023.

| Image Name | IoU | f1 | Kappa | Precision | Recall | Accuracy | Area ($Km^2$) |
|---|---|---|---|---|---|---|---|
| Formentera_3_august_2023 | 93.78 | 96.77 | 93.76 | 96.78 | 96.77 | 96.77 | 29.04 |
| Sur_Oeste_Menorca_9_august_2023 | 93.71 | 96.55 | 83.18 | 96.65 | 96.7 | 96.7 | 16.4 |
| PortoColom_CalaMillor_13_july_2023 | 92.21 | 95.85 | 91.41 | 95.9 | 95.89 | 95.89 | 28.23 |
| Sur_Ibiza_26_august_2023 | 90.07 | 94.56 | 86.7 | 94.62 | 94.69 | 94.69 | 22.55 |
| North_Cabrera_14_july_2023 | 87.8 | 92.87 | 69.54 | 93.3 | 93.28 | 93.28 | 1.36 |
| Pollença_27_june_2023 | 87.68 | 93.0 | 77.37 | 93.16 | 93.34 | 93.34 | 26.05 |
| Capdepera_31_july_2023 | 87.3 | 92.93 | 88.4 | 93.2 | 93.17 | 93.17 | 15.88 |
| South_Cabrera_29_june_2023 | 86.38 | 92.42 | 81.93 | 92.38 | 92.5 | 92.5 | 1.53 |
| Sur_Este_Menorca_9_august_2023 | 85.02 | 91.69 | 87.11 | 92.36 | 91.93 | 91.93 | 16.57 |
| Norte_Ibiza_27_june_2023 | 84.32 | 91.28 | 87.09 | 91.95 | 91.45 | 91.45 | 4.42 |
| Alcudia_15_july_2023 | 81.08 | 87.67 | 50.55 | 90.25 | 89.27 | 89.27 | 76.2 |
| Oeste_Ibiza_19_july_2023 | 80.96 | 89.21 | 80.97 | 90.04 | 89.59 | 89.59 | 7.74 |
| Es_Trenc_15_july_2023 | 79.47 | 86.24 | 67.01 | 91.98 | 86.47 | 86.47 | 12.66 |
| Este_Ibiza_14_july_2023 | 78.24 | 87.03 | 69.24 | 87.84 | 87.69 | 87.69 | 13.27 |
| CalaPi_CalaFiguera_12_july_2023 | 76.17 | 85.29 | 69.68 | 87.56 | 86.37 | 86.37 | 36.52 |
| Banyalbufar_Soller_12_july_2023 | 75.25 | 84.9 | 72.54 | 87.29 | 86.13 | 86.13 | 8.85 |
| Dragonera_Banyalbufar_16_july_2023 | 72.61 | 83.72 | 73.37 | 86.49 | 83.65 | 83.65 | 7.17 |
| Palma_29_june_2023 | 68.32 | 80.13 | 63.45 | 81.64 | 81.39 | 81.39 | 62.94 |
| Norte_Menorca_23_june_2023 | 67.65 | 79.55 | 69.89 | 86.56 | 78.33 | 78.33 | 14.16 |
| Este_Menorca_23_june_2023 | 66.3 | 79.37 | 69.65 | 83.97 | 79.37 | 79.37 | 27.04 |
| Soller_Calobra_18_july_2023 | 44.43 | 60.2 | 40.03 | 66.68 | 60.43 | 60.43 | 6.0 |

**Table 11:** Averge performance metrics for the final model with 2023 images.

| Method | IoU | f1 | Kappa | Precision | Recall | Accuracy |
|---|---|---|---|---|---|---|
| voting_top_10 | 79.92 | 87.63 | 71.69 | 89.46 | 88.26 | 88.26 |
| voting_top_5 | 79.38 | 87.24 | 71.14 | 88.95 | 87.87 | 87.87 |
| resnet34 | 79.29 | 87.28 | 71.34 | 88.56 | 87.71 | 87.71 |
| voting_top_smart | 79.29 | 87.2 | 70.49 | 89.13 | 87.83 | 87.83 |
| voting_top_3 | 79.27 | 87.11 | 70.95 | 88.88 | 87.74 | 87.74 |
| inceptionresnetv2 | 79.17 | 87.2 | 70.96 | 88.42 | 87.75 | 87.75 |
| mobilenetv2 | 77.76 | 86.34 | 69.05 | 87.46 | 86.84 | 86.84 |
| resnext101 | 77.61 | 85.9 | 68.89 | 87.7 | 86.58 | 86.58 |
| seresnet152 | 77.31 | 86.01 | 68.99 | 87.31 | 86.51 | 86.51 |
| inceptionv3 | 77 | 85.71 | 68.53 | 87.22 | 86.13 | 86.13 |
| resnet152 | 76.81 | 85.43 | 67.79 | 87.3 | 86.07 | 86.07 |
| seresnext101 | 76.52 | 85.23 | 67.32 | 86.95 | 85.98 | 85.98 |
| efficientnetb7 | 75.22 | 84.42 | 65.69 | 85.85 | 85.18 | 85.18 |
| densenet201 | 74.85 | 84.18 | 65.63 | 86.19 | 84.63 | 84.63 |
